# Matrimeres are systemic nanoscale mediators of tissue integrity and function

**DOI:** 10.1101/2024.03.25.586585

**Authors:** Koushik Debnath, Irfan Qayoom, Steven O’Donnell, Julia Ekiert, Can Wang, Mark A. Sanborn, Chang Liu, Ambar Rivera, Ik Sung Cho, Saiumamaheswari Saichellappa, Peter T. Toth, Dolly Mehta, Jalees Rehman, Xiaoping Du, Yu Gao, Jae-Won Shin

## Abstract

Tissue barriers must be rapidly restored after injury to promote regeneration. However, the mechanism behind this process is unclear, particularly in cases where the underlying extracellular matrix is still compromised. Here, we report the discovery of matrimeres as constitutive nanoscale mediators of tissue integrity and function. We define matrimeres as non-vesicular nanoparticles secreted by cells, distinguished by a primary composition comprising at least one matrix protein and DNA molecules serving as scaffolds. Mesenchymal stromal cells assemble matrimeres from fibronectin and DNA within acidic intracellular compartments. Drawing inspiration from this biological process, we have achieved the successful reconstitution of matrimeres without cells. This was accomplished by using purified matrix proteins, including fibronectin and vitronectin, and DNA molecules under optimal acidic pH conditions, guided by the heparin-binding domain and phosphate backbone, respectively. Plasma fibronectin matrimeres circulate in the blood at homeostasis but exhibit a 10-fold decrease during systemic inflammatory injury *in vivo*. Exogenous matrimeres rapidly restore vascular integrity by actively reannealing endothelial cells post-injury and remain persistent in the host tissue matrix. The scalable production of matrimeres holds promise as a biologically inspired platform for regenerative nanomedicine.

---

The extracellular matrix (ECM) is the anchor for many types of cells and harbors various biophysical and biochemical signals essential to maintain the function of these cells including the barrier function of vascular endothelial cells^1^. The ECM controls how cells generate physical forces and tension via integrin-mediated mechanotransduction^2^. Cell-ECM interactions via integrins is essential for establishment and stabilization of endothelial barriers^3,4^. The disruption of adherens junctions represents a unifying pathological feature of various inflammatory and traumatic injuries, which can contribute to persistent tissue damage and fibrosis^5^. Genetically downregulating the key components of endothelial cell-ECM mechanotransduction, including β_1_ ^6^ and β_3_ ^7^ integrins that bind to Arg-Gly-Asp (RGD), and focal adhesion kinase (FAK)^8,9^ spontaneously results in edema. However, it remains unclear how specific ECM components constitutively maintain vascular integrity and tissue function.

Fibronectin (FN) is an ECM glycoprotein abundant in plasma^10^. FN plays an important role in tissue repair and regeneration, since it can be rapidly deposited at the site of blood vessel injury^11^ independently of platelets^12^. The activation of integrin-FAK in endothelial cells requires an insoluble form of FN^13^. It has been long appreciated that fibrillar networks of FN are assembled *de novo* when soluble FN binds to integrin on the plasma membrane^14^ in the presence of cell-generated physical forces^15^ *in vitro*. Whether this process is responsible for maintaining and restoring vascular integrity remains unclear. Recent studies suggest that cell-secreted nanoscale mediators transport ECM molecules, including FN^16–19^. However, nanoscale mediators consist of heterogeneous subpopulations, including lipid membrane-bound extracellular vesicles (EVs) and non-vesicular extracellular particles (NVEPs)^20^. It is unclear which subpopulation specifically bears the molecules and properties needed to competently activate ECM signaling over distance. Here, we report the discovery of matrimeres, which are non-vesicular nanoparticles assembled by the interaction between ECM proteins and DNA molecules, and act as endogenous nanoscale mediators that safeguard against tissue injury.

## Results

### Quantification of cell-secreted FN^+^ NVEPs

Mesenchymal stromal cells (MSCs) are localized in the perivascular niche and act as tissue architects by producing various paracrine mediators to orchestrate the host response^21^. MSCs can also package cargo molecules into EVs and secrete them^22–24^. MSCs are significant sources of cellular FN in different tissues^25^. Thus, we initially sought to test the presence of FN in EVs from MSCs. Clonally derived mouse bone marrow D1 MSCs were used as previously described, since clonal populations provide less cell-to-cell heterogeneity than primary cells^23,24^. Conventional differential centrifugation was used to obtain a crude pellet of small (< 200 nm) nanoscale mediators from the medium of cultured MSCs. Given the heterogeneity of nanoscale mediators secreted by cells^26–29^, we further used a gel-based immunoaffinity approach combined with nanoparticle tracking analysis to determine the fraction of the pellet that expresses the ECM protein FN of the cellular origin (i.e., mouse-specific) or the tetraspanin protein CD63 that is present on the EV membrane. The pellet consists of 55.1% CD63^+^, 46.5% FN^+^ subpopulations (**Fig. 1A, i**). Human primary bone marrow MSCs also secrete a substantial fraction (∼30%) of FN^+^ subpopulations (**Fig. S1A**). About 98% of the total fraction is pulled down by using both CD63 and FN antibodies simultaneously (**Fig. 1A, i**), suggesting that there is a minimum overlap between CD63 and FN subpopulations (∼3%). Biochemical disruption by Triton-X (TX) (**Fig. S1B**) was then performed to determine the fraction of lipid-containing structures^30^ in the pellet after negative selection. FN depletion makes the nanostructures in the pellet more sensitive to TX-mediated disruption, while CD63 depletion or simultaneous FN and CD63 depletion makes them more resistant (**Fig. 1A, ii; Fig. S1C**). Based on these results, it is possible to calculate 8 nanoscale subpopulations in terms of CD63 expression, FN expression, and TX sensitivity (**Supplementary Text**). The majority of MSC-secreted subpopulations consist of TX-resistant FN^+^CD63^-^(37.9%) and TX-sensitive FN^-^CD63^+^ (41.8%) (**Fig. 1A, iii**).

**Figure 1.**
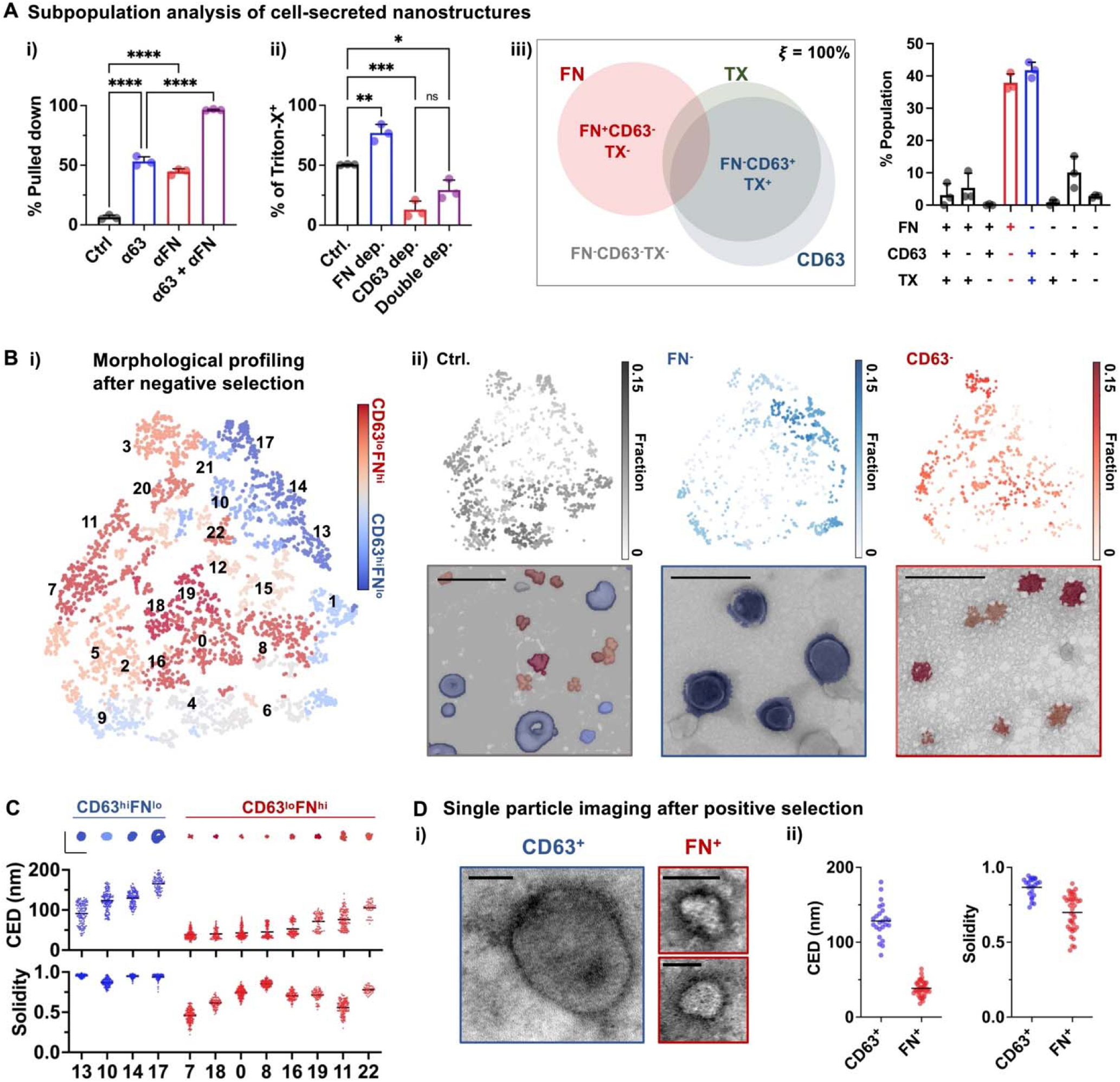
FN^+^ nanoscale subpopulations secreted from MSCs are NVEPs. **(A)** The nanostructure subpopulations from mouse D1 MSCs. (i) Percentage of subpopulations pulled down by anti-mouse CD63 (αCD63), anti-mouse FN (αFN), or both, compared to biotin-only control (ctrl.) determined by the gel-based pull-down assay. (ii) Percentage of the triton-X (TX) sensitive fraction before (ctrl.) and after depletion (dep.) with FN or CD63 antibody. \**p* < 0.05, \*\**p* < 0.01, \*\*\**p* < 0.001, \*\*\*\**p* < 0.0001 via one-way ANOVA with Tukey’s multiple comparisons test. (iii) Quantification of nanoscale subpopulations in terms of CD63, FN, and TX sensitivity. (Left) Proportional Venn diagram. (Right) Percentage of subpopulations. *n* = 3 experiments. **(B)** Morphological profiling of small nanostructures from mouse D1 MSCs after negative selection with FN or CD63 antibody-coated magnetic beads. (i) T-distributed stochastic neighbor embedding (t-SNE) plot based on 12 morphological parameters of small nanostructures from 9-11 TEM images of crude (ctrl., *n* = 1581), FN-depleted (*n* = 895), and CD63-depleted (*n* = 679) samples pooled from 3 different batches. Cluster numbers are assigned by the Leiden clustering algorithm. The color scheme is based on the relative representation of each nanostructure cluster by CD63-depleted (CD63^lo^, hence FN^hi^, red) *vs.* FN-depleted (FN^lo^, hence CD63^hi^, blue) samples. (ii) Fraction of clusters represented by nanostructures in each group. (Above) Visualization of cluster fractions under the t-SNE plot. (Below) Representative TEM images from each group with an overlay from the color scheme in (i). Scale bar = 300 nm. **(C)** Quantification of morphological parameters for each nanostructure cluster after negative selection. (Above) Representative nanostructures. Scale bar = 200 nm. (Below) Circular equivalent diameter (CED) and solidity. *n* = 47-198 for each group. **(D)** Confirmation of nanostructures after positive selection with FN or CD63 antibody and visualization of single vesicles or particles. (i) Representative images showing vesicular CD63^+^ and non-vesicular FN^+^ nanostructures. Scale bar = 50 nm. (ii) CED and solidity. *n* = 24 for CD63^+^ and *n* = 38 for FN^+^ nanostructures. The error bars denote s.d.

Quantitative analysis of transmission electron microscopy (TEM) images obtained after negative selection by a magnetic bead-based pull-down approach (**Fig. S1D**) identifies multiple nanoscale subpopulations (**Fig. S2A**). Each subpopulation was classified based on the relative representation by FN-depleted *vs.* CD63-depleted fractions—*i.e.,* CD63^lo^FN^hi^ *vs.* CD63^hi^FN^lo^ (**Fig. 1B**). The non-vesicular CD63^lo^FN^hi^ subpopulations are generally smaller (40 ∼ 105 nm) than the vesicular CD63^hi^FN^lo^ subpopulations (90 ∼ 165 nm) with varying degrees of solidity (**Fig. 1C**) and other morphological features (**Fig. S2B**). The CD63^lo^FN^hi^ subpopulations morphologically resemble previously described NVEPs, such as exomeres^26^ and supermeres^29^ rather than EVs. TEM analysis of the positively selected fractions after pulling down with the antibodies attached to small (∼8 nm in diameter; **Fig. S3A**) magnetic nanoparticles confirms that single FN^+^ particles are non-vesicular (**Fig. 1D**). Thus, MSCs secrete a substantial fraction of FN^+^ NVEPs distinct from EVs.

### Cell-secreted FN^+^ NVEPs maintain endothelial integrity after inflammatory injury

Both FN^+^ NVEPs and CD63^+^ EVs are present in the plasma of mouse peripheral blood *in vivo* (**Fig. 2A, i**). There are ∼4.0 x 10^6^ nanostructures per μl of mouse blood plasma (**Fig. 2A, ii, iii**), and ∼20% of them express FN (**Fig. 2A, iv**). Given that the blood volume of a mouse is ∼80 μl/g^31^, the results suggest that there are ∼1.3 x 10^9^ plasma FN^+^ NVEPs in blood per 20g mouse (**Fig. 2A, v**). Human blood plasma also contains a large fraction (∼50%) of FN^+^ NVEPs (**Fig. S3B**). Systemic treatment of mice with lipopolysaccharide (LPS) - the endotoxin that is widely used to disrupt vascular integrity, and hence the host ECM^32^ and endothelial junctions^33^ - decreases the fraction of FN^+^ NVEPs, resulting in the reduced number of FN^+^ NVEPs in blood plasma by ∼10-fold within 8 h (**Fig. 2A**). Mass spectrometry analysis was performed to define the proteomes of FN^+^ NVEPs. The proteomics datasets were acquired first using data-dependent acquisition mode to generate a spectral library and then with data-independent acquisition mode for identification and quantitation, all proteomics data were searched with ProLuCID^34^ and MSFragger^35^, filtered at 1% protein false discovery rate, visualized by the Sequence Coverage Visualizer^36^, and compared to the MatrisomeDB reference database^37,38^. We found that Fibronectin 1 (Fn1) was the most abundant protein in both plasma (∼26.0%) and MSC-secreted (∼56.6%) FN^+^ NVEPs (**Fig. 2B, i**; **Supplementary Data S1**). Both FN^+^ NVEPs contain a substantial fraction of albumin (Alb) and immunoglobulin heavy constant mu (Ighm) (5∼6%). Plasma FN^+^ NVEPs contains more fibrinogen proteins (e.g., Fgg, Fgb, Fga) and pregnancy zone protein (Pzp), while MSC-secreted FN^+^ NVEPs contain more actin beta (Actb), fetuin-A (Ahsg), and glyceraldehyde 3-phosphate dehydrogenase (Gapdh) proteins. There is diversity in isoforms of immunoglobulin and complement proteins between plasma (C1qc, Jchain, C1qa, C1qb) and MSC-secreted (Igg3, C3, Cp, C4b) FN^+^ NVEPs. Fn1 is similar in both FN^+^ NVEPs in terms of sequence coverage (∼70% total) with some differences in post-translational modifications towards the N-terminus (**Fig. 2B, ii; Fig. S4**). Several domains known to be important in FN function were detected in Fn1 of both FN^+^ NVEPs, including a cell attachment domain (Arg-Gly-Asp; RGD, 1614-1616), three heparin binding domains (HBDs), and collagen binding domain. The EIIIB domain was not detected, the EIIA domain was partially detected (plasma: 30.7%, MSC: 44.3% coverage; **Supplementary Data S2**), and most of the V domain were detected in Fn1 of both FN^+^ NVEPs. This suggests that MSCs likely assemble FN^+^ NVEPs from the Fn1 variant known to be abundant in plasma^39^.

**Figure 2.**
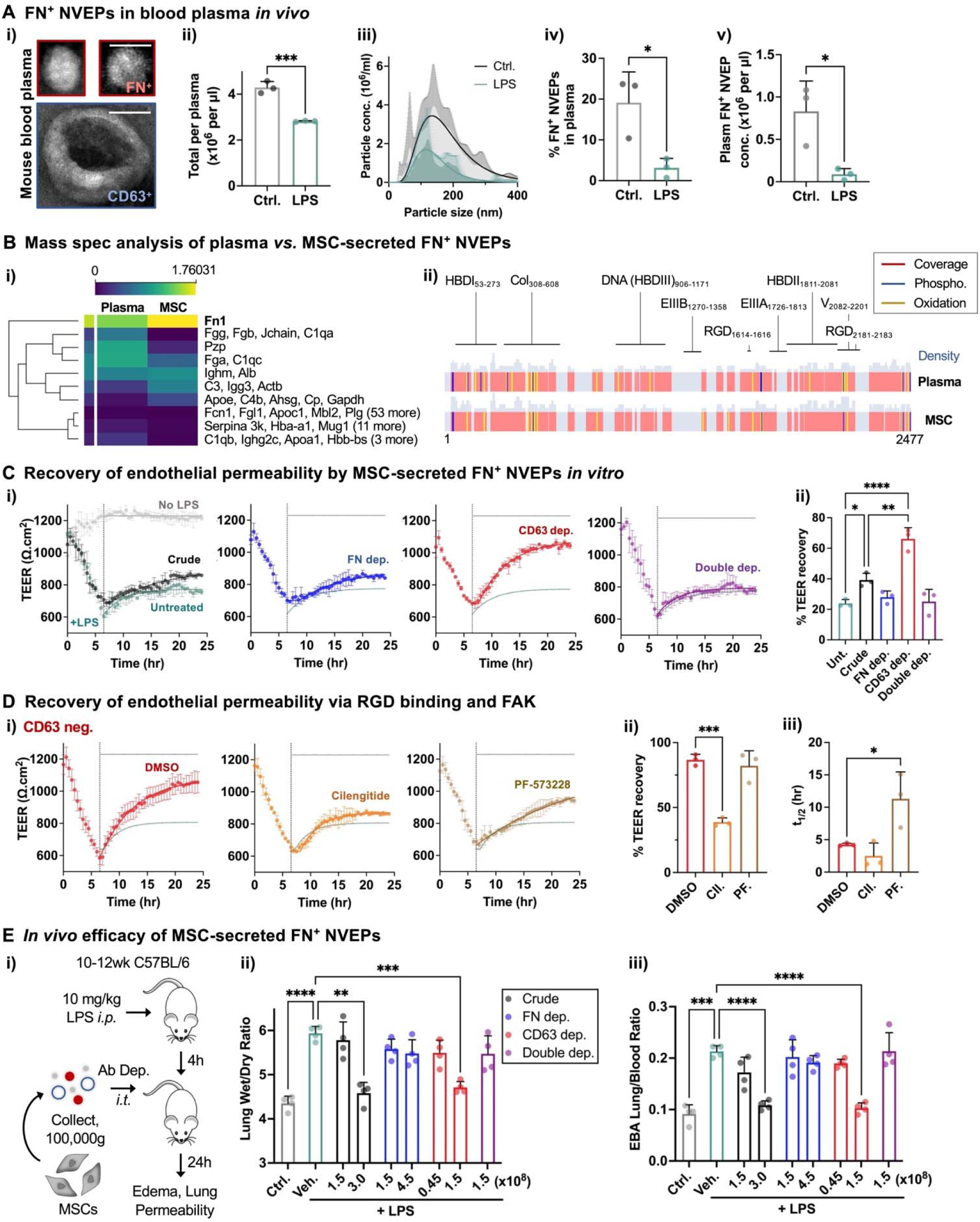
FN^+^ NVEPs from MSCs are enriched with the activity to restore endothelial barrier function. **(A)** FN^+^ NVEPs are present in mouse blood plasma. (i) Representative TEM images showing the distinct morphology of FN^+^ NVEPs *vs.* CD63^+^ EVs in plasma after positive selection with the respective antibodies. Scale bar = 50 nm. (ii) Concentration of nanostructures. (iii) Hydrodynamic sizes of nanostructures. The average data points were fitted to log-normal distribution curves. (iv) Percentage of FN^+^ matrimeres. (v) Concentration of FN^+^ NVEPs. For (ii)-(v), mice were also injected intraperitoneally (*i.p.*) with lipopolysaccharide (LPS, 10 mg/kg), and blood plasma were analyzed after 8 h. *n* = 3 mice, \**p* < 0.05, \*\*\**p* < 0.001 via unpaired T-test. **(B)** Mass spectrometry analysis of FN^+^ NVEPs from MSCs *vs.* plasma. (i) Proteome coverage comparison; label-free protein quantitation based on log-transformed intensity. (ii) Fn1 sequence coverage comparison. Red denotes detected regions; histograms above denote peptide overlap density; blue and yellow denote the phosphorylation and oxidation sites detected, respectively. Pooled from *n* = 2 biological replicates. **(C)** Effects of fractionated small nanostructures from mouse D1 MSCs on restoration of endothelial barrier function after LPS treatment *in vitro*. The HUVEC monolayer plated on TEER electrodes was treated with LPS (1 μM) for 6 h, followed by washout and addition of 1.5 x 10^8^ small nanostructures per well after gel-based depletion (dep.) with FN and/or CD63 antibody at *t* = 6.5 h. (i) TEER kinetics over 24 h. The y-axis starts from 500 Ω.cm^2^. A dotted vertical line indicates the time when the samples were added. The data points from *t* = 6.5 h to 24 h were fitted to one-phase association curves. (ii) Percentage of TEER recovery after LPS treatment from *t* = 6.5 h to 24 h. *n* = 3 experiments. **(D)** Effects of inhibiting RGD and FAK on the recovery of TEER by the CD63-depleted fraction after LPS treatment. DMSO, cilengitide (200 nM) or PF-573228 (100 nM) was added simultaneously with CD63^-^nanostructures. (i) TEER kinetics. (ii) Percentage of TEER recovery. (iii) Half-time (*t_1/2_*) of TEER recovery. *n* = 3 experiments. **(E)** Effects of fractionated small nanostructures from MSCs on restoration of lung endothelial barrier function after acute lung injury *in vivo*. (i) Overview of strategy to determine the efficacy of fractionated nanostructures from MSCs in a mouse model of LPS-induced lung injury. Mice were treated with LPS (10 mg/kg) for 4 h, followed by intratracheal (*i.t.*) administration of mouse D1 MSC-secreted nanostructures after antibody (ab)-based depletion (dep.). (ii) Lung edema by quantifying lung tissue wet-dry ratio. (iii) Lung vascular permeability by quantifying Evans blue albumin (EBA) accumulation. The doses are indicated in the unit of 10^8^ per 20g mouse. *n* = 4 mice for each group. \**p* < 0.05, \*\**p* < 0.01, \*\*\**p* < 0.001, \*\*\*\**p* < 0.0001 via one-way ANOVA with Tukey’s multiple comparisons test for **(C)** and **(D)**, and via Welch’s one-way ANOVA with Dunnett’s T3 post-test for **(E)**. The error bars denote s.d.

Based on these studies, we tested the hypothesis that providing exogeneous MSC-secreted FN^+^ NVEPs accelerates the resolution of tissue injury by restoring endothelial barrier integrity. The trans-endothelial electrical resistance (TEER) assay was used to test the role of FN^+^ matrimeres from MSCs in restoring the integrity and permeability of the endothelial cell monolayer *in vitro*. LPS decreases the TEER value over 6 h, and washing out LPS alone results in ∼20% of TEER recovery in 24 h (**Fig. 2C**). Adding 1.5 x 10^8^ of the crude nanoscale fraction from MSCs increases TEER recovery to ∼40%, while the CD63-depleted fraction enhances TEER recovery to over 60% (**Fig. 2C**). In contrast, neither FN-depleted nor CD63 and FN double-depleted fraction was able to rescue TEER at this dose (**Fig. 2C**). Thus, FN^+^ NVEPs from MSCs are enriched with the ability to directly reverse LPS-induced endothelial hyperpermeability *in vitro*. Competitive inhibition of integrin binding by cilengitide (cyclic RGD) decreases the extent of TEER recovery, while inhibition of focal adhesion kinase (FAK) activity by PF-573228 delays the kinetics of TEER recovery by the CD63-depleted fraction (**Fig. 2D**). Thus, restoration of endothelial barrier integrity by FN^+^ NVEPs requires integrin ligation, and may involve FAK-mediated integrin signaling. We then tested whether FN^+^ NVEPs from MSCs can reverse endotoxemia-induced inflammatory injury in mice. After 4 h of intraperitoneal (*i.p.*) LPS treatment, MSC-secreted nanoscale fractions were delivered to mice via the intratracheal (*i.t.*) route in order to localize the delivery to the lungs (**Fig. 2E, i**). Without fractionation, 3.0 x 10^8^ particles per 20g mouse are necessary to reverse LPS-induced lung edema and vascular permeability 24 h after delivery (**Fig. 2E, ii, iii**). In contrast, the FN-depleted fraction does not show the effects even at a higher dose, 4.5 x 10^8^/20g (**Fig. 2E, ii, iii**). On the other hand, CD63 depletion (or FN enrichment) reduces the effective dose to 1.5 x 10^8^/20g, while removing FN^+^ fraction after CD63 depletion abolishes the therapeutic effects (**Fig. 2E, ii, iii**). Since 37.9% of MSC-secreted nanoscale fractions are TX-resistant FN^+^CD63^-^particles (**Fig. 1)**, ∼1.14 x 10^8^ MSC-secreted FN^+^ NVEPs per 20g mouse are needed to restore endothelial barrier function in the lungs after LPS injury.

### Matrimeres are NVEPs assembled from ECM proteins and DNA molecules

We next assessed how FN^+^ NVEPs are formed by MSCs. DNA is known to be present in both EVs and NVEPs^26–29^. A significantly higher amount of DNA is present in the CD63-depleted fraction than the FN-depleted fraction (**Fig. 3A, i**). Treatment of MSC-secreted nanoscale fractions with DNase I (50 U/ml) *ex vivo* dramatically decreases the number of the FN^+^ fraction but not that of CD63^+^ fraction (**Fig. 3A, ii**). Pre-treatment of the CD63-depleted fraction with DNase I abolishes its therapeutic effects in LPS-induced lung injury *in vivo* (**Fig. 3B**). Thus, DNA is essential for the formation of functional FN^+^ NVEPs. Here, we define matrimeres as NVEPs that are assembled from at least one ECM protein as the major protein component and DNA molecules as scaffolds.

**Figure 3.**
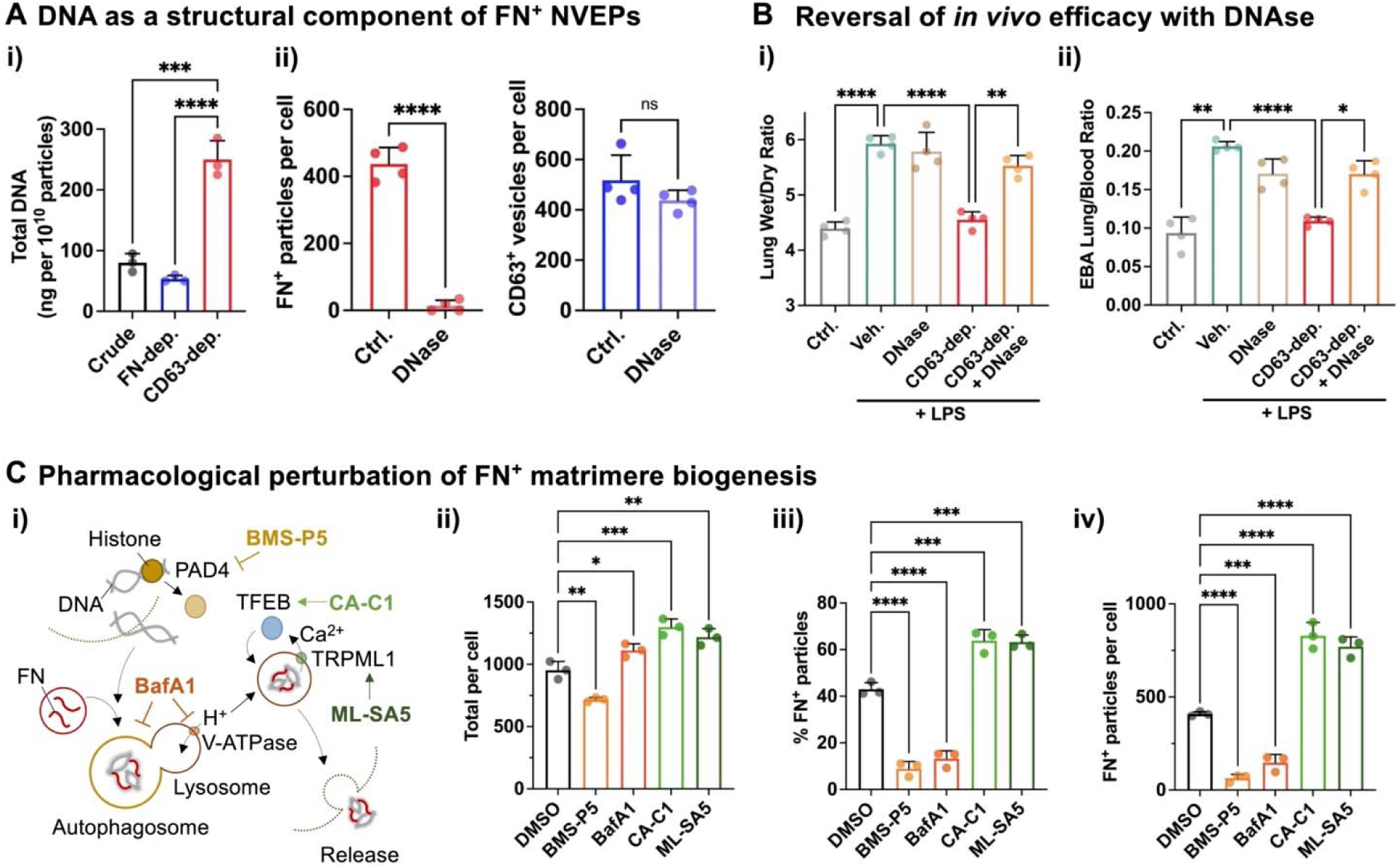
DNA is essential for cells to form FN^+^ matrimeres. **(A)** Evaluation of DNA as a structural component of FN^+^ matrimeres. (i) Total amount of DNA in mouse D1 MSC-secreted nanostructures without depletion (crude) or after gel-based depletion (dep.) with FN or CD63 antibody. (ii) Sensitivity of (left) FN^+^ matrimeres and (right) CD63^+^ EVs to DNase I treatment *ex vivo*. The crude pellet from MSCs was treated without (ctrl.) or with DNase I (50 U/ml) for 1 h at 37°C, followed by washout and pull-down assay to determine the number of FN^+^ matrimeres or CD63^+^ EVs normalized by the number of MSCs. \*\*\*\**p*<0.0001, ns: not significant via unpaired T-test. *n* = 4 experiments. **(B)** Effects of DNase I on the efficacy of the CD63-depleted fraction (1.5 x 10^8^ per 20g mouse) in LPS-induced lung injury *in vivo*. (i) Lung tissue wet-dry ratio. (iii) EBA accumulation in lung tissue. *n* = 4 mice for each group. **(C)** Effects of pharmacological agents on FN^+^ matrimere production from MSCs. (i) Overview of the tested drugs and their known targets. Mouse D1 MSCs were cultured in the presence of DMSO, BMS-P5 (1 μM), BafA1 (200 nM), CA-C1 (1 μM), or ML-SA5 (20 μM) for 1 day, followed by quantification of FN^+^ matrimeres secreted per cell. (ii) Number of small nanostructures per cell. (iii) Percentage of FN^+^ matrimeres. (iv) Number of FN^+^ matrimeres per cell. *n* = 3 experiments. \**p* < 0.05, \*\**p* < 0.01, \*\*\**p* < 0.001, \*\*\*\**p* < 0.0001 via one-way ANOVA with Tukey’s multiple comparisons test for (A), i and (D) ii-iv, and via Welch’s one-way ANOVA with Dunnett’s T3 post-test for (B). The error bars denote s.d.

We next studies potential intracellular pathways which drive FN^+^ matrimere biogenesis by MSCs (**Fig. 3C, i**). For excess DNA to be released from the nucleus, arginine in histone undergoes modification from a positively charged ketimine group to a neutrally charged ketone group via citrullination, which is mediated by peptidylarginine deiminase (PAD)^40^. BMS-P5, a selective inhibitor against PAD4^41^ decreases the number FN^+^ matrimeres secreted from MSCs (**Fig. 3C, ii-iv**). Bafilomycin-A1 (BafA1), which blocks both autophagosome-lysosome fusion and lysosome acidification via vacuolar H^+^ ATPase^42^ also decreases the number of MSC-secreted FN^+^ matrimeres (**Fig. 3C, ii-iv**). Conversely, activating the components of lysosome function either by ML-SA5 (agonist of transient receptor potential mucolipin 1, or TRPML1)^43^ or curcumin analog C1 (CA-C1, agonist of transcription factor EB, or TFEB)^44^ significantly increases FN^+^ particle number per cell (**Fig. 3C, ii-iv**). Most MSCs (>90%) remained viable after drug treatment (**Fig. S5**). Thus, the assembly of FN and DNA into matrimeres depends on acidic intracellular compartments.

### In vitro reconstitution of matrimeres

Unlike EVs where cargo molecules are protected by the lipid bilayer, the formation, and stability of matrimeres will likely rely more on direct interactions among constituent molecules. These interactions can potentially occur in an acidic environment (**Fig. 3D**). FN contains several heparin-binding domains (HBDs) (**Fig. 2B, ii**), which consist of arginine and lysine residues^45^. Arginine contains guanidinium cation, which forms hydrogen bonds with negatively charged phosphate, sulfate, and carboxylate groups^46^. In particular, the guanidinium group is known to interact more strongly with phosphate than with sulfate or carboxylate due to a larger difference in electronegativity^47^. Supporting this notion, earlier studies show that FN can directly bind to DNA that contains a phosphate backbone^48^ via a HBDIII^49^ (**Fig. 2B, ii**). However, the relevance of these studies to nanoscale assembly remains unclear. Thus, we tested whether matrimeres can be reconstituted *in vitro* in acidic pH (**Fig. 4A, i**). Affinity-purified mouse plasma FN (10 μg/ml) was incubated with sonicated DNA molecules (10 μg/ml) from mouse MSCs with an equal w/v ratio. The maximum number of reconstituted plasma FN-DNA matrimeres (∼1.5 x 10^10^/ml) were formed at pH 5.5, as opposed to other tested pH values (**Fig. 4A, ii**). The zeta potential of the reconstituted FN-DNA matrimeres remains close to zero at pH 7.4, while it becomes slightly positive at pH 4.5 (**Fig. 4A, iii**), consistent with the feature of zwitterionic nanoparticles. On average, 1.5 x 10^10^ FN-DNA matrimeres contain ∼4.6 μg of plasma FN (**Fig. 4A, iv**), and ∼5 μg of DNA (**Fig. 4A, v**), suggesting that the conversion efficiency from the input materials to FN-DNA matrimeres is ∼50%. Thus, ∼840 FN monomers and ∼307,700 DNA base pairs are incorporated into a single particle (**Supplementary Text**). Purified mouse plasma FN samples and assembled FN-DNA matrimeres show the similar proteome composition with Fn1 as the most abundant protein (86∼90%), and the similar post-translational modifications of Fn1 (**Fig. S6; Supplementary Data S3 and S4**). DNA size profiling confirms that the sonification process fragments the DNA isolated from MSCs, while purified FN interacts with a broad range (250-10000 bp; peak ∼ 1500 bp) of DNA molecules to form FN-DNA matrimeres (**Fig. 4B**). We next addressed whether FN-DNA matrimeres are selectively enriched in specific DNA regions. We observed that sequencing coverage depth was generally similar between DNA assembled into reconstituted FN-DNA matrimeres and genomic DNA (**Fig. S7**), suggesting that there is no intrinsic enrichment of specific DNA sequences to form nanoscale complexes with FN. TEM analysis shows that reconstituted plasma FN-DNA matrimeres exhibit NVEP-like structures (**Fig. 4C, i**) with the mean size ∼33 nm and varying degrees of solidity (**Fig. 4C, ii**). Incubating the recombinant C-terminal HBD from FN (‘HBDII’; human FNIII_12-13_, 181 a.a., 91% homology to mouse) with DNA molecules is sufficient to form nanoparticles at pH 5.5 (**Fig. 4D**), suggesting that HBD is important in the assembly of FN-DNA matrimeres. Incubating purified FN with the anionic inorganic polymer, long-chain polyphosphate (‘polyP’, p700) also results in nanoparticle formation (**Fig. 4D**), confirming the notion that phosphate backbone is an essential building block of reconstituted matrimeres. Vitronectin (VTN) from mouse plasma, another ECM glycoprotein that contains RGD and an HBD^50^, also forms nanoparticles with DNA molecules (**Fig. 4D**). After assembly and resuspension in PBS at pH 7.4, the particle number of FN-DNA and VTN-DNA matrimeres remains constant over 9 days; in contrast, the number of FN-polyP decreases by 50% with *t_1/2_* ∼ 4 h but remains constant afterward (**Fig. 4E**), the phenotype that could be attributed to rapid hydrolysis of polyP^51^ as opposed to DNA molecules. The results show a facile approach to reconstitute a catalog of matrimeres based on biomolecular interactions between ECM proteins and DNA molecules at an optimal acidic pH similar to that of late endosomes and lysosome compartments^52,53^.

**Figure 4.**
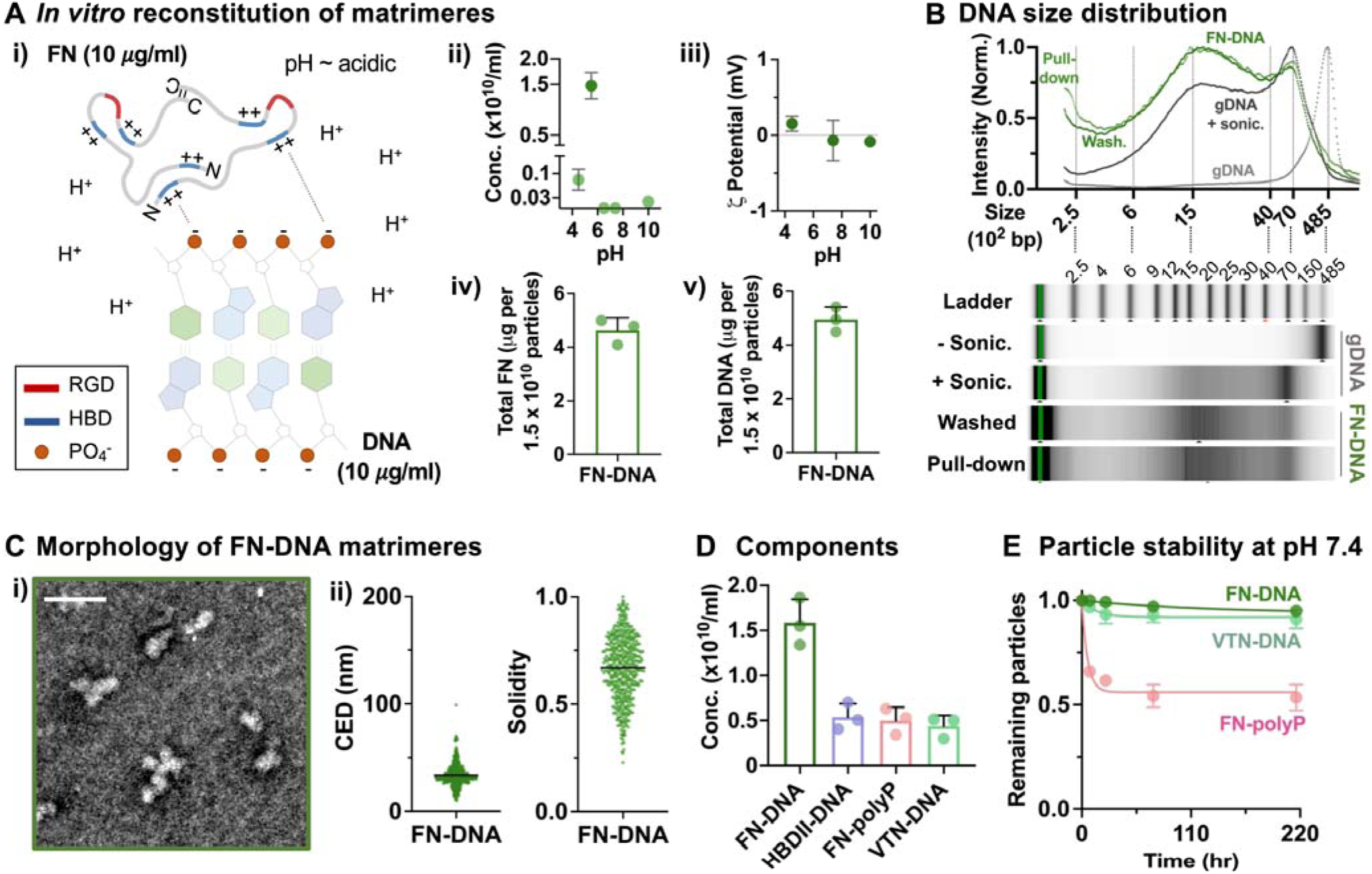
Matrimeres can be reconstituted *in vitro* based on biomolecular interactions between ECM and DNA molecules. **(A)** Reconstitution of matrimeres *in vitro*. (i) Schematic showing the putative biomolecular interactions between heparin binding domain (HBD) of FN proteins, which are often present as dimers, and the phosphate backbone of DNA molecules under acidic pH. Matrimeres were reconstituted from purified FN protein and sonicated genomic DNA molecules isolated from mouse D1 MSCs in a non-phosphate, acetate buffer at 4°C overnight. (ii) Effects of pH on *in vitro* reconstitution of FN-DNA matrimeres measured right after formation. (iii) Zeta potential of reconstituted FN-DNA matrimeres at different pH values. (iv) Quantification of FN protein incorporated into FN-DNA matrimeres. (v) Quantification of DNA incorporated into FN-DNA matrimeres. *n* = 3 batches. **(B)** Size distribution of DNA molecules in FN-DNA matrimeres. (Above) Quantification of DNA size distribution from genomic DNA (gDNA) without sonication, gDNA with sonication (sonic.), FN-DNA matrimeres after washout, and FN-DNA matrimeres after pull-down with FN antibody. (Below) Representative images from DNA electrophoresis. **(C)** TEM analysis of FN-DNA matrimeres. (i) Representative image. Scale bar = 50 nm. (ii) Quantification of (Left) CED and (Right) solidity. *n* = 501 particles from 10 images pooled from 3 different batches. **(D)** Formation of matrimeres from different molecular components at pH = 5.5. All the components were mixed at 10 μg/ml in a 1:1 ratio. *n* = 3 batches for each group. **(E)** Stability of reconstituted matrimeres at neutral pH *in vitro*. The data points were fitted to one-phase decay curves. *t_1/2_* and plateau for FN-DNA: 0.94, 89.4 h; VTN-DNA: 0.92, 10.3 h; FN-polyP: 0.46, 4.0 h. *n* = 3 batches for each group. The error bars denote s.d.

### Reconstituted matrimeres remain in tissue after delivery

We sought to test the biodistribution of reconstituted matrimeres in mice after delivery into the lungs. Animal imaging of fluorescently conjugated, reconstituted plasma FN-DNA matrimeres in mice (**Fig. S8**) shows that right after *i.t.* delivery, 68.7% of the signals are present in the lungs, while some signals can also be detected in the liver (22.5%) and the kidneys (7.8%). After 8 hours, ∼50% of matrimeres in the lungs are cleared, but the remaining signals are persistent over 9 days in the lungs, while the signals are no longer detected in other organs (**Fig. 5A**). To further characterize the continued residence of fluorescent FN-DNA matrimeres in the lungs at a higher resolution, we performed three-dimensional (3D) two-photon imaging analysis of fresh lung samples^54^. There are ∼5 large, cell-like fluorescent particles per ∼530,000 μm^3^ lung tissue after both 1 and 9 days of delivery (**Fig. 5B, i, ii**). Supporting this observation, tissue-resident alveolar macrophages show the highest signals from fluorescent FN-DNA matrimeres among the tested immune cell subpopulations in the lungs (**Fig. S9**) after 1 day of delivery (**Fig. 5B, iii**). Thus, macrophages are expected to clear the nanoparticles^55^. However, the results show that ∼10% of both vasculature (CD31^+^) and collagen fiber surfaces are contacted by the signals from fluorescent FN-DNA matrimeres after 1 day of delivery. In comparison, this number increases to ∼30% for collagen fibers after 9 days of delivery due to increased contact by fluorescent FN-DNA matrimeres over time (**Fig. 5C**). Thus, plasma FN is deposited in the lung parenchyma after delivery of reconstituted FN-DNA matrimeres.

**Figure 5.**
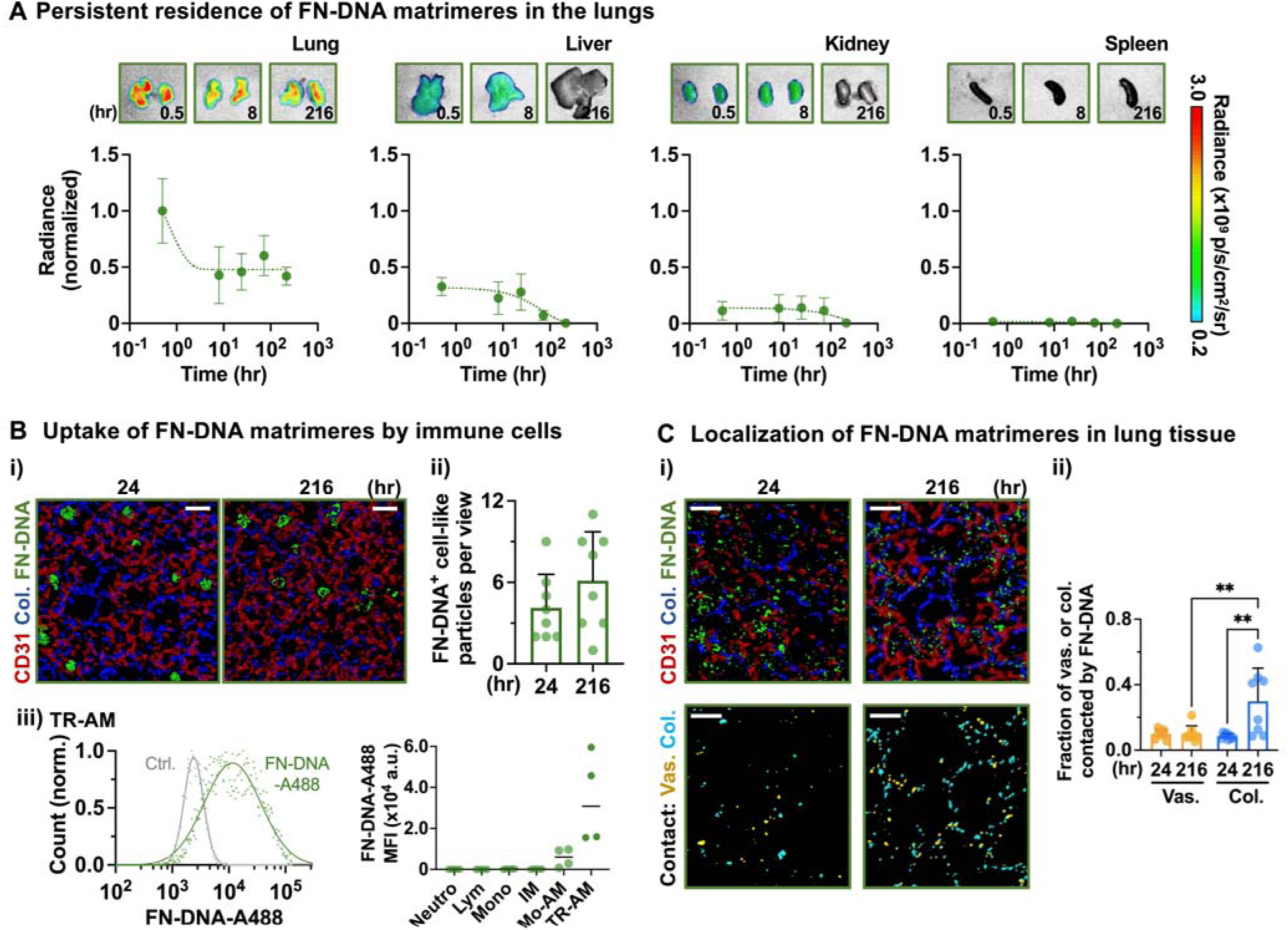
Reconstituted FN-DNA matrimeres persist in tissue after delivery. **(A)** Biodistribution of FN-DNA matrimeres in different organs. Mice were treated LPS (7.5 mg/kg) for 4 h, followed by delivery of Cy7-conjugated FN-DNA matrimeres via the *i.t.* route and IVIS imaging at different time points. Radiance values from Cy7 signals were normalized to the mean value of the data points from *t* = 0.5 h in lung tissue. The data points were fitted to one-phase decay curves. *t_1/2_* and plateau for lung: 2.1h, 0.47; liver: 42.4h, 0; kidney: 110.4h, 0. *n* = 4 mice for each time point. **(B)** Uptake of FN-DNA matrimeres by lung tissue-resident cells. Two-photon analysis was done in isolated lung tissue after *i.t.* delivery of AZDye 488 (A488)-conjugated FN-DNA matrimeres (green) into LPS-treated mice. Prior to delivery, mice were injected with the Alexa Fluor 647-conjugated antibody against CD31 (red) to visualize the vasculature, while collagen fibers (blue) were visualized by second harmonic generation. (i) Representative images from two-photon microscopy showing cell-like A488^+^ particles (>7 μm in circular equivalent diameter) in lung tissue. Scale bar = 20 μm. (ii) Quantification of cell-like A488^+^ particles per field of view. *n* = 8 tissue regions from 4 mice for each time point. (iii) Flow cytometry analysis of A488^+^ signals in neutrophils (Neutro), lymphocytes (Lym), monocytes (Mono), monocyte-derived alveolar macrophages (Mo-AM), tissue-resident alveolar macrophages (TR-AM), and interstitial macrophages (IM). (Left) Representative histogram showing A488^+^ signals in TR-AMs (*vs*. negative control). (Right) Quantification of A488^+^ mean fluorescent intensity (MFI) in each immune subpopulation. *n* = 4 mice. **(C)** Localization of FN-DNA matrimeres in lung tissue. (i) Representative images showing (above) small A488^+^ clusters in lung tissue after delivery, and (below) the vasculature (yellow) and the collagen (cyan) areas contacted by A488^+^ signals. Scale bar = 20 μm. (ii) Quantification of FN-DNA localization in lung tissue in terms of the fraction of the vasculature or the collagen surface areas contacted by A488^+^ signals per field of view. *n* = 8 tissue regions from 4 mice for each time point. \*\**p* < 0.01 via Welch’s one-way ANOVA with Dunnett’s T3 post-test. The error bars denote s.d.

### Reconstituted matrimeres restore endothelial integrity and tissue function after inflammatory injury

We tested the therapeutic efficacy of reconstituted matrimeres in mice after inflammatory tissue injury. Adding 1.5 x 10^8^ reconstituted plasma FN-DNA matrimeres rapidly rescues TEER of the endothelial monolayer after LPS treatment with *t_1/2_* ∼ 1.0 h in an RGD-dependent manner (**Fig. 6A**), which is faster than the CD63-depleted (or FN-enriched) fraction from MSCs (**Fig. 2D, iii**) at the same dose. FN-DNA matrimeres successfully reverse LPS-induced lung edema and vascular permeability 24 h after delivery in a dose-dependent manner—a lower dose (0.75 x 10^8^) is still effective in treating lung injury (**Fig. 6B**). In contrast, neither instillation of soluble FN nor HBDII-DNA particles can rescue vascular integrity after LPS treatment (**Fig. 6B**). FN-polyP particles show the therapeutic effects but require a higher dose (4.5 x 10^8^), correlating with the rapid degradation of this formulation at pH 7.4 *in vitro* (**Fig. 4E**). The therapeutic effects can also be achieved with VTN-DNA matrimeres albeit with a higher dose (2.5 x 10^8^; **Fig. 6B**), suggesting the generalizability of the approach to other RGD and HBD-containing ECM proteins. We observed improved lung function by FN-DNA matrimeres after injury as shown by restoration of inspiratory capacity, lung tissue damping, compliance and elastance to uninjured levels (**Fig. 6C**). To investigate whether the results can be generalized to sterile inflammatory injury, we also tested the efficacy of reconstituted plasma FN-DNA matrimeres following cutaneous reverse passive Arthus (rpA) reaction, which models immune complex-mediated, type III hypersensitivity injury^56^ (**Fig. 6D, i**). The intravenous (*i.v.*) injection of FN-DNA matrimeres (3.0 x 10^8^) is sufficient to inhibit not only rpA-induced vascular permeability (**Fig. 6D, ii**) but also bleeding (**Fig. 6D, iii**), whereas soluble FN does not show the effect. Together, reconstituted matrimeres are sufficient to effectively restore vascular integrity and tissue function after inflammatory injury *in vivo*.

**Figure 6.**
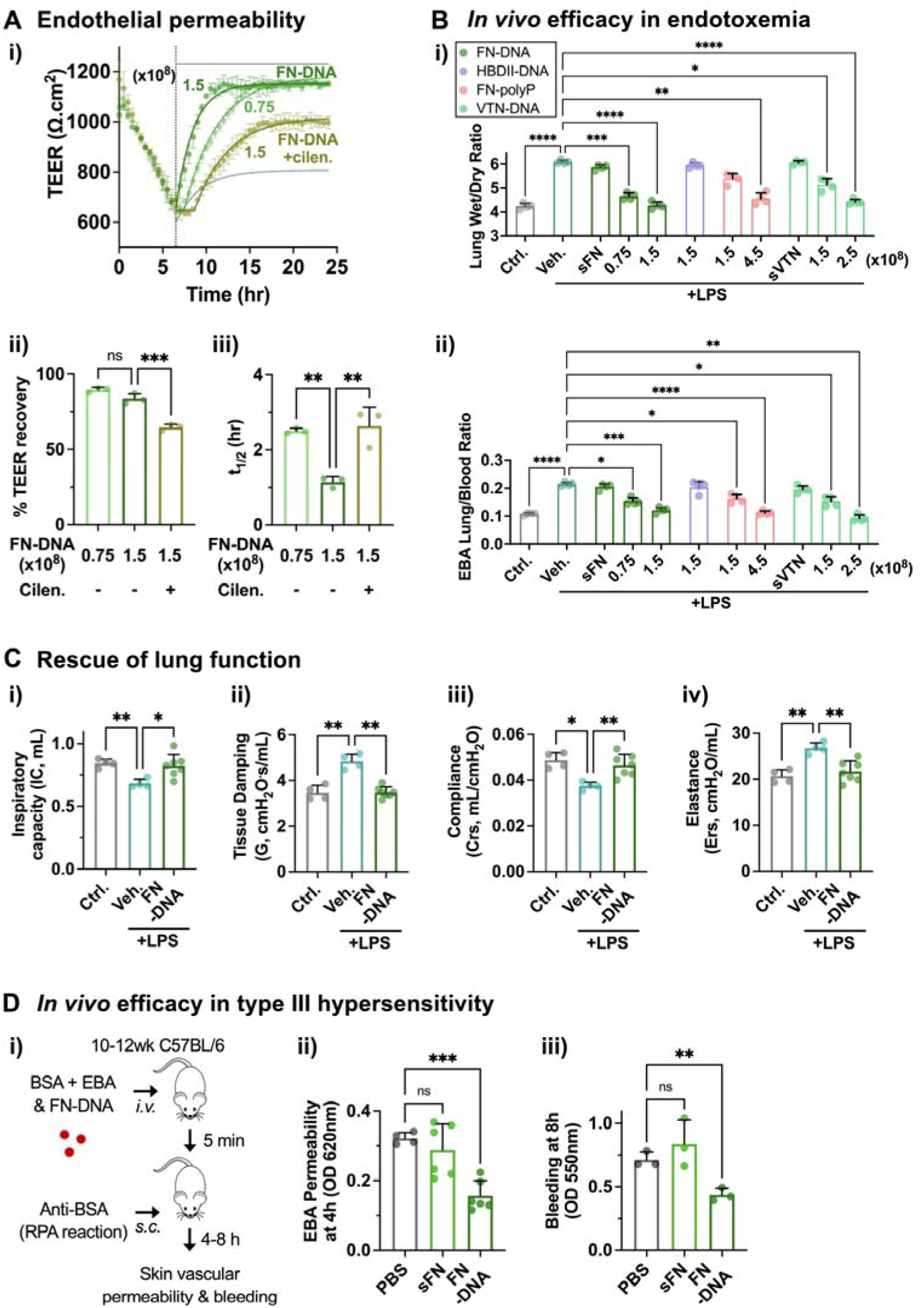
Reconstituted matrimeres restore vascular integrity after inflammatory injury. **(A)** Effects of reconstituted FN-DNA matrimeres on the recovery of TEER after 6 h LPS (1 μM) treatment *in vitro*. (i) TEER kinetics. (ii) Percentage of TEER recovery. (iii) Half-life (*t_1/2_*) of TEER recovery. *n* = 3 experiments. **(B)** Effects of reconstituted matrimeres on restoration of lung endothelial barrier function after LPS (10 mg/kg)-induced lung injury *in vivo*. (i) Lung tissue wet-dry ratio. (iii) EBA accumulation in lung tissue. Soluble FN (sFN) and VTN (sVTN) protein groups (10 μg/ml, 30 μl per 20g mouse) were included as controls. The doses are indicated in the unit of 10^8^ per 20g mouse. *n* = 4 mice for each group. **(C)** Effects of FN-DNA matrimeres (1.5 x 10^8^ per 20g mouse) on restoring lung function after LPS-induced injury *in vivo* in terms of (i) inspiratory capacity, (ii) tissue damping, (iii) compliance, and (iv) elastance. *n* = 4 mice for control and LPS + vehicle and *n* = 7 mice for LPS + FN-DNA. **(D)** Effects of FN-DNA matrimeres on restoration of skin endothelial barrier function after cutaneous reverse passive Arthus (rpA) reaction *in vivo*. (i) Overview of rpA model. Mice were intravenously (*i.v.*) injected with BSA and either PBS, soluble FN (100 ng/20g) or FN-DNA matrimeres (3.0 x 10^8^/20g). EBA was injected to evaluate vascular permeability. After 5 min, anti-BSA was injected subcutaneously to induce rpA reaction, followed by analysis of skin (ii) EBA permeability after 4 h or (iii) bleeding after 8 h. For (ii), *n* = 4 mice for PBS and *n* = 6 mice for soluble FN and FN-DNA matrimeres. For (iii), *n* = 3 mice for each group. \**p* < 0.05, \*\**p* < 0.01, \*\*\**p* < 0.001, \*\*\*\**p* < 0.0001 via one-way ANOVA with Tukey’s multiple comparisons test for **(A)**, and via Welch’s one-way ANOVA with Dunnett’s T3 post-test for **(B)**-**(D)**. The error bars denote s.d.

## Discussion

Our results suggest that cells employ a mechanism to constitutively maintain tissue integrity by assembling and systemically circulating nanoscale matrimeres. We initially anticipated that small EVs would play a role in delivering ECM molecules. However, a number of studies that reported FN on EVs performed no further purification step after initial ultracentrifugation^17–19^. In one study, ultracentrifugation via iodixanol density gradient was used to show the presence of FN in the fraction that also expresses EV membrane markers^16^. However, FN was found to be of bovine origin from serum used in cell culture, rather than of cellular origin; the relative abundance of bovine FN^+^ EVs compared to other nanoscale fractions is still unclear. Interestingly, the same study and another one^27^ also show the presence of FN in higher density, non-EV fractions. A recent study analyzed cell-secreted crude nanoscale fractions from large-scale production after pre-treating them with DNase to show that heparin affinity chromatography enriches for both FN^+^ EVs and NVEPs^57^. However, it is unclear from this study how many DNase-resistant FN^+^ fractions are secreted per cell. By quantitative analysis, our work presents the surprising finding that most of cell-secreted FN^+^ nanoscale mediators are non-vesicular, detergent resistant, and require DNA to form.

Matrimeres are distinct from macromolecules formed by direct assembly between FN molecules without DNA, which requires forced FN unfolding by cells^15^ and results in fibril formation. In contrast, DNA enables the packaging of multiple ECM molecules into a single matrimere at high density without fibril formation. Soluble plasma FN is globular in a dimerized form with 8∼10 nm in hydrodynamic radius^58,59^. This size is smaller than the known persistence length of double-stranded DNA in a physiological salt solution (∼50 nm or 150 base pairs)^60^. Thus, the natural variation in size and sphericity of matrimeres is likely due to varied length of incorporated DNA molecules. This nanoparticle design by nature offers multivalent ligand presentation necessary to constitutively activate ECM signaling compared to soluble circulating ECM molecules. In addition, matrimeres offer an effective mechanism to systemically circulate a payload of ECM molecules that can be deployed to restore damaged tissue.

A large quantity of soluble plasma FN (∼200 μg/ml in mouse)^10^ is secreted from hepatocytes^61^. However, our results show that soluble FN alone is not sufficient to restore endothelial junctions after inflammatory injuries *in vivo*. In contrast, we show that as low as 0.75 x 10^8^/20g reconstituted FN-DNA matrimeres, which contain only 25 ng plasma FN, restore endothelial junctions when the total number of circulating plasma FN^+^ matrimeres goes down to 1.3 x 10^8^/20g after systemic treatment with LPS. In addition to lung injury, endotoxemia induces hepatocyte apoptosis, which is exacerbated by liver-specific knockout of plasma FN^62^. Hepatocytes are also known to contain numerous acidic intracellular vesicles^63^ and release extracellular DNA^64^. Thus, hepatocytes may contribute to circulating plasma FN^+^ matrimeres. However, it is possible that other cells, such as MSCs can also contribute to plasma FN^+^ matrimeres given their similarity in Fn1 sequence. Cell-specific genetic deletion studies could define the relative contribution of each cell type to the plasma FN^+^ matrimere turnover. Since EIIIA and EIIIB domains are normally deficient in plasma FN but become included after inflammatory injury^65,66^, it is possible that the presence of these alternative exons interferes with FN^+^ matrimere formation, thereby reducing the assembly of circulating FN^+^ matrimeres upon inflammatory injury. The estimated effective dose of MSC-secreted FN^+^ matrimeres is ∼50% higher than that of reconstituted plasma FN-DNA matrimeres. This can be due to other proteins that may sterically hinder RGD presentation or post-translational modifications specifically present or absent in Fn1 of MSC-secreted matrimeres.

Conventionally, extracellular DNA is thought to be released by dying or cancerous cells, and hence serves as a danger signal to induce inflammatory response as exemplified by neutrophil extracellular traps^67^. However, a small amount (a ng/ml range) of DNA is normally present in peripheral blood^68^. In addition, lymphocytes^69^ and fibroblasts^70^ secrete DNA-protein complexes in the absence of stimulation. While DNA is known to be present in exomeres and supermeres^26,29^, our results show that DNA molecules act as scaffolds to constitutively assemble and release matrimeres for maintenance of barrier function. It is important to note that matrimeres are distinct from circulating DNA that forms complexes with histones^71^, since histone was neither detectable in matrimeres nor necessary to form matrimeres. However, it is possible that changes in epigenetic modifications under disease conditions can influence chromatin accessibility, and hence matrimere turnover and DNA composition. Deeper, long-read DNA sequencing analysis^72^ could provide further insights into the genetic and epigenetic composition of matrimeres in health and disease. While the autophagosome-lysosome machinery normally degrades cytoplasmic chromatin fragments and proteins^73^, we show that acidity promotes interaction between ECM proteins and DNA molecules, which are subsequently secreted as matrimeres. The biogenesis mechanism of FN^+^ matrimeres is distinct from EV biogenesis since contrary to what we observed with FN^+^ matrimeres, inhibition of autophagosome-lysosome fusion is known to enhance CD63^+^ EV secretion^74,75^. From a functional perspective, this process supports an emerging role of secretory autophagy or lysosomal exocytosis in tissue repair and regeneration as previously shown in the context of plasma membrane repair and bone resorption^76^.

MSC-secreted crude nanoscale fractions were previously shown to resolve preclinical models of acute tissue injuries^77,78^, and some of these efforts have already advanced to clinical trials (e.g., NCT05354141). In this context, EV cargo transfer has been proposed as a main mechanism for the resolution of acute tissue injuries^77,78^. However, our study presents quantitative evidence that FN^+^ matrimeres from MSCs rather than EVs are enriched with therapeutic activity to resolve LPS-induced lung injury and can directly restore endothelial integrity by activating integrin signaling. Furthermore, our work provides a viable path for clinical translation by enabling the synthesis of functional matrimeres without a need to use living cells with broad potential to treat systemic and local inflammation. A deeper understanding of ECM-DNA interactions powered by biophysical and machine learning approaches will inform how to manufacture matrimeres from purely synthetic peptides and nucleic acids with precisely defined shape, properties, and function. Since our data show that plasma FN^+^ matrimeres are significantly decreased upon inflammatory injury, FN^+^ matrimeres also have potential as mechanistically direct biomarkers to screen for the severity of patients with inflammatory injuries who can then be treated by exogenous matrimeres. In summary, this study opens new avenues of investigations into biogenesis, structure, and function of matrimeres, presenting broad opportunities to design biologically-inspired regenerative nanostructures.

## Methods

### Cell Culture

All cells were cultured in an incubator at 37°C and 5% CO_2_. Mouse bone marrow D1 mesenchymal stromal cells (MSCs; #CRL-12424, ATCC) were cultured using high-glucose Dulbecco’s Modified Eagle Medium (DMEM; Thermo) supplemented with 10% v/v premium-grade fetal bovine serum (FBS; #S11550, Atlanta Biologicals), 100 μg/ml penicillin/streptomycin (P/S; Thermo), and 2 mM GlutaMAX (Thermo) to 80% confluency before passaging (up to passage number 25). Human primary bone marrow MSCs derived by plastic adherence of mononucleated cells from human bone marrow aspirate donors (Lonza) and were cultured in α-minimal essential medium (αMEM; Thermo) supplemented with 20% v/v FBS, 100 μg/ml P/S, and 2 mM GlutaMAX to 80% confluency before passaging (up to passage number 6). Human umbilical vein endothelial cells (HUVEC; #CC-2519, Lonza) were cultured using Endothelial Cell Growth Basal Medium-2 (EBM-2; #CC-3156, Lonza) supplemented with 10% FBS and 1% antibiotic-antimycotic and endothelial cell growth supplement (#CC-4176, Lonza) to 80% confluency before passaging (up to passage number 8). All cells were cultured at 37°C in 5% CO_2_ and routinely tested for mycoplasma contamination.

### Nanostructure size and number characterization

Nanoparticle Tracking Analysis 3.2 (NTA) via NanoSight NS300 (Malvern) using a 405 nm laser was used to obtain particle size and number. Samples were introduced by syringe pump at a rate 100 μL/min. Three 30 s videos were acquired using camera level 16 followed by detection threshold 7. The software automatically set the camera focus, shutter, blur, minimum track length, minimum expected particle size, and maximum jump length. Samples were diluted as needed to maintain particles per video from 100 to 2,000. All samples were tested as compared to appropriate blank conditions in order to ensure specificity. For some experiments, samples were treated with 0.1% (v/v) Triton X-100 (Sigma) for 10 min at room temperature (RT) prior to analysis.

### Differential centrifugation to obtain crude small nanostructure fractions from cultured cells

Cells were first cultured in 175 mm^2^ using high-glucose DMEM supplemented with 10% FBS, 1% P/S, and 1% GlutaMAX to 80% confluency. Cells were then washed once with Hanks’ Balanced Salt Solution (HBSS; Thermo) followed by incubation with serum-free DMEM for 1 h. Afterward, the medium was changed to DMEM with 5% exosome-depleted FBS (Thermo). After 48 h, the medium was centrifuged at 2,000g for 10 min to remove cell debris followed by centrifugation at 10,000g to remove large EVs (>200 nm). The supernatant was then added slowly to a 14-mL polystyrene ultracentrifuge tube (Beckman) containing 1.5 mL of 30% sucrose (Thermo) in phosphate-buffered saline (PBS; Thermo), followed by centrifugation at 100,000g for 90 min. The upper non-sucrose layer was aspirated and washed with PBS followed by another centrifugation at 100,000g for 90 min. The crude pellet consisting of small (< 200 nm) nanostructures was resuspended in PBS and analyzed by NanoSight NS300 prior to downstream experiments.

### Isolation of crude nanostructure fractions from blood plasma

C57BL6/J mice (#000664, Jackson Lab) were injected intraperitoneally (*i.p.*) with PBS or 10 mg/kg of lipopolysaccharide (LPS from *E. coli* O111:B4, 1 µM; #L2630, Sigma). After 8 h, blood was drawn (500∼600 µL per mouse) by cardiac puncture and collected in a 1.5-mL centrifuge tube containing 10 µL of 5 mM EDTA solution as an anticoagulant. Blood plasma was collected by centrifugation at 3,000 g for 15 min to obtain the supernatant, followed by another round of centrifugation at 3,000g for 20 min. Fresh human blood plasma was purchased from ZenBio (#SER-PLE10ML-SDS, anticoagulant: EDTA). The plasma was then diluted at 1:20 with ice-cold PBS, followed by the same differential centrifugation procedure as the samples from cultured cells.

### Immunoaffinity-based negative selection with antibody-functionalized magnetic beads

Streptavidin-coated magnetic beads (Dynabeads Biotin Binder; #11047, Thermo) were washed 3 times with PBS followed by incubation with biotin anti-mouse CD63 antibody (Clone REA563; Miltenyi Biotec) or biotin anti-mouse fibronectin (FN) antibody (#IRBAMSFBNGFBL; Innovative Research). 4.0 x 10^8^ streptavidin-coated magnetic beads were used to incubate with 20 µg of the biotinylated antibody either at RT for 1 h or at 4 °C for 8 h with gentle tilting and rotating. The beads were then washed 3 times with PBS and pull-down with a magnet to remove unbound antibody molecules. Typically, 1.0 x 10^7^ to 10^8^ antibody-attached beads were incubated with a 10-times higher number of small nanostructures at RT for 1 h with gentle tilting and rotating. After incubation, the beads were removed by a magnet, followed by centrifugation at 1000 g for 2 min. The supernatant was collected for downstream analyses.

### Immunoaffinity-based negative selection with antibody-functionalized hydrogels

1 mg of streptavidin (Agilent) was dissolved in 370 µL of deionized (DI) water, and 30 µL of 7.5% Na_2_CO_3_ (890 µM) was added to the solution to make the pH ∼ 8. The solution was kept at 4 °C with gentle stirring for 10 min followed by adding 100 µl of acrylic acid N-hydroxysuccinimide ester (1.35 mg/mL freshly prepared in Milli-Q water). The reaction was continued for 4 h, followed by adding 50 µL poly(ethylene glycol)-diacrylate (PEG-DA; Sigma) solution (density: 1.12 mg/mL) and 50 µl of lithium phenyl-2,4,6-trimethyl-benzoyl phosphinate (LAP; Sigma) solution (5 mg/mL freshly prepared in Milli-Q water). Afterward, 900 µL of the solution was placed into a 6-well plate, followed by the exposure of the solution to 365 nm ultraviolet light for 1 min to polymerize the solution into the gel. The gel was washed 3 times with PBS to remove excess streptavidin and LAP. The gel was incubated with biotin anti-mouse CD63 antibody or biotin anti-mouse FN antibody at 1 µg/mL final concentration for 1 h at RT with gentle tilting and shaking. After incubation, the gel was washed with PBS three times to remove unbound antibodies. Each gel was used to deplete a specific population by incubating with 1.0 x 10^8^ to 10^9^ small nanostructures for 1 h at RT, and the supernatant was collected for downstream experiments.

### Immunoaffinity-based positive selection by antibody-conjugated small magnetic nanoparticles

Fe (II)-stearate was used as a precursor to synthesize hydrophobic magnetic iron-oxide (γ-Fe_2_O_3_) nanoparticles via a high-temperature organometallic approach. 4.267 g of stearic acid (#175366; Sigma) was dissolved in methanol at ∼50 °C, followed by adding 6.136 mL of 25% tetramethylammonium hydroxide solution (#331635, Sigma) dropwise, and stirring for 12 h. Once the solution became transparent, freshly prepared methanolic FeCl_3_ (#8.03945; Sigma) solution (0.811 g FeCl_3_ in 2 mL methanol) was added dropwise with vigorous stirring. Stirring was continued for another 30 min until the yellow precipitate turned into a brownish color. The precipitate was then collected via filtration and washed with warm methanol several times for complete removal of free stearic acid. Next, 0.161 g of octadecyl amine (#8.41029, Sigma) and 0.161 g of 4-methylmorpholine N-oxide (#224286, Sigma) were dissolved in 12 mL 1-octadecene (#O806, Sigma) in a three-necked round bottom flask. Then, 0.313 g of Fe (II)-stearate powder was added to the solution. The solution was heated up to 300 °C temperature under argon gas flow for 10 min. Afterward, the solution was cooled down to RT instantly by purging liquid nitrogen. The colloidal solution of hydrophobic magnetic iron-oxide nanoparticles (∼8 nm in size; **Fig. S3A**) was further purified by precipitating in excess ethanol and redissolving in cyclohexane (Alfa Aesar). The nanoparticles and acrylate monomers were dissolved into reverse micelles (cyclohexane: IGEPAL: water = 15: 4: 1), and polyacrylate formation was initiated under an argon atmosphere in the presence of persulfate (Sigma) for 15 min. N-(3-Aminopropyl)methacrylamide (APMA) hydrochloride (Sigma) was used to introduce the amine functional group. About 400 to 500 primary amine groups were present per polymer-coated nanoparticle on the surface, and were used for conjugation with dibenzocyclooctyne (DBCO)-N-hydroxysuccinimidyl (NHS) ester (#A133, Click Chemistry Tools). Typically, ∼5.0 x 10^13^ of the nanoparticles were added to DI water, and 3.3 mg of DBCO-NHS in 100 µL DMSO solution was added dropwise with vigorous stirring. The reaction took place for 4 h, followed by removal of free DBCO via magnetic pull-down. In parallel, 50 ng/μl of purified anti-mouse CD63 antibody (Clone 446703, R&D Systems) or purified anti-mouse FN antibody (#IRBAMSFBNGF; Innovative Research) was conjugated to 0.3 mM azidoacetic acid NHS ester (#1070, Click Chemistry Tools) in a 300 μl PBS. The DBCO-functionalized magnetic nanoparticles (5.0 x 10^11^) and azide-functionalized antibodies (15 μg) were mixed in PBS and gently agitated for 2 h at 4 °C for copper-free click chemistry. The antibody-functionalized nanoparticles (2.0 x 10^11^) were washed out 3 times with PBS for complete removal of unconjugated antibodies, and reacted with nanostructures (1.0 x 10^10^) for positive selection, followed by downstream analyses.

### Negative-stain transmission electron microscopy (TEM)

10 μL of samples with an optimized concentration (1.0∼1.5 x 10^10^ /100 μL range) in double-filtered PBS solution were deposited onto the TEM grid (Formvar/Carbon 300 Mesh, Copper; #FCF300-CU, Electron Microscopy Sciences) and incubated for 10 min. Excess liquid was blotted by the Whatman 42 filter paper, followed by negative staining with 8 μL of 6% uranyl acetate (SPI Supplies) in Milli-Q water for 20 s at RT in dark. Excess liquid was blotted, followed by aerial drying in dark for 10 min. Grids were examined via JEOL JEM-1400F TEM, operating at 80 kV. Digital micrographs were acquired using the AMT-BIOSPRINT12M-B (mid-mount) and AMT software (Version 701).

### Morphological analysis of nanostructures from TEM images

The images from negative-stain TEM were acquired under the same condition (Exposure: 960 ms x 2 std. frames, gain: 1, bin: 1) for computational analysis of nanostructure morphology. Each image was processed by automatic background subtraction (rolling ball radius: 50), noise removal by despeckle, and contrast enhancement with saturated pixel at 0.3% by ImageJ (NIH). Image threshold was set at top 10-20% of the intensity histogram. The density-based spatial clustering of applications with noise (DBSCAN) algorithm was then used to identify and digitize individual nanostructures^79^ in each image via a custom Python (3.11) program. The algorithm groups together localizations that are within a radius (*ε*), given a minimum number of localizations to be considered a cluster (*ρ*). Since each pixel ∼ 1 nm, the *ε* value was set at 5, and the *ρ* value was set at 35. Large aggregates (>200 nm) were excluded from the analysis. The shape of each nanostructure was quantified by the following morphological parameters: area, circular equivalent diameter (CED), perimeter, convex area, convex perimeter, solidity, convexity, circularity, roundness, least bounding rectangle length (LBRL), least bounding rectangle width (LBRW), and aspect ratio. The coordinate points consisting of each nanostructure were also recorded to map the analyzed values back to the original images.

### Morphological classification of cell-secreted nanostructure subpopulations from TEM images after negative selection

To classify nanostructures based on multiple morphological parameters, dimension reduction and clustering techniques were implemented by the Scanpy toolkit (1.9) in Python. The data in each morphological parameter were normalized by the StandardScaler module, followed by putting the normalized data in an annotated matrix (AnnData package). The data from control samples were put into a reference matrix, while the data from the samples after negative selection with CD63 or FN antibody were put into another matrix. The batch correction between two matrices was done by implementing the batch balanced k nearest neighbors (BBKNN) algorithm^80^, followed by combining them into one matrix. The Leiden algorithm^81^ was then used to cluster nanostructures into multiple groups. The relative representation of each cluster by data points from CD63-depleted *vs.* FN-depleted samples was calculated and used for color coding. The clustered data were visualized by t-distributed stochastic neighbor embedding (t-SNE)^82^, which is a commonly used dimensionality reduction technique.

### Measurement of endothelial monolayer permeability by trans-endothelial electrical resistance (TEER) assay

HUVECs were seeded at 40,000 cells per well on gold-plated electrodes (8W10E PET; Applied Biosciences) and cultured in complete medium for 24 h until a confluent monolayer was formed as indicated by the TEER value of ∼1,100 Ω.cm^2^. The cells were then washed with HBSS followed by adding basal medium without or with LPS. TEER was recorded every 30 min using the Electric Cell-Substrate Impedance Sensing (ECIS) system at 37°C and 5% CO_2_ (Applied Biophysics). LPS was washed away after 6 h, followed by adding an indicated number of nanostructures. For some experiments, cilengitide (200 nM, # 22289) or PF-573228 (100 nM, #14924) (both from Cayman Chemical) was added together with the nanostructures. TEER was further recorded until 24 h. The data points between 6.5 and 24 h were fitted to standard one-phase association curves to calculate plateau and half-life (t_1/2_) values. The percentage of TEER recovery was calculated by taking the difference between TEER values at *t* = 24 h and 6.5 h for each experiment, and divide it by the average difference between TEER values from the control without LPS and LPS at *t* = 6.5 h.

### Animal model of endotoxemia-induced lung injury

The animal procedures were done in accordance with NIH and institutional guidelines approved by the ethical committee from the University of Illinois at Chicago. C57BL/6J mice at age of 8∼10 weeks were treated with LPS (10 mg/kg) *i.p.* to induce acute lung injury. After 4 h, mice were anesthetized by ketamine/xylazine (50/5 mg/kg) *i.p.*, and the samples were administered by intratracheal (*i.t.*) instillation at an indicated dose. One day after LPS administration, mice were evaluated for lung edema and vascular permeability as described previously^83^. Mice were anesthetized using ketamine/xylazine (50/5 mg/kg) *i.p.* and a solution of Evans blue albumin (EBA) (20 mg/kg) was applied via retro-orbital intravenous (*i.v.*) injection. After 40 min, mice were sacrificed, and lung tissue was harvested along with a fraction of circulating blood. Right lung tissue was weighed initially (wet weight) and after 36 h of incubation at 65 °C (dry weight) to calculate the wet/dry ratio. Left lung tissue was homogenized, and Evans blue was extracted with formamide. Evans blue content was measured by absorbance at 620 nm and normalized to that present in circulating blood.

### Preparation of peptides for LC-MS/MS analysis

Samples were prepared in the following manner: female C57BL6/J mouse blood plasma in PBS, pH 7.4, was spun down at 18,000 x g, 4°C for 15 minutes to ensure removal of cell debris. The supernatant was transferred to a 1.5 mL Eppendorf lo-bind tube. Blood plasma was then sonicated at 30%, at 55 kHz for thirty seconds, three times with 15 second intervals on ice in between to aid membrane protein solubilization. Protein was quantified with BCA Protein Assay (Pierce, Appleton, WI, cat. A55864). After pulling down the samples with mouse fibronectin antibody-functionalized Fe_2_O_3_ particles (1∼2 µg fibronectin), they were resuspended in PBS, pH 8.5 and CaCl_2_ (1mM final), were taken for overnight digestion with mass spec grade Trypsin/Lys-C Mix (Promega, Madison, WI, cat. No. V5071, 1:20). Samples were reduced with TCEP (final concentration 5 mM) and subsequently alkylated with 2-Chloroaceetamide (final concentration 50 mM). Samples were digested overnight (16 h) at 37°C. Digestion was stopped with the addition of formic acid (5% final).

### Proteomics data acquisition

Total digests (equivalent of 1-2 µg) were prepared for LC-MS/MS analysis with Evotips (Evosep, Odense, Denmark, cat. EV2011) for subsequent desalting and loading steps. Samples were injected for LC-MS/MS analysis with Q Exactive HF mass spectrometer (ThermoFisher, Waltham, MA) coupled with Evosep One (Evosep, Odense, Denmark) standardized liquid chromatography platform. Peptides were separated by a capillary C18 reverse-phase column (Evosep, Odense, Denmark, Endurance, 15 cm x 150 µm, 1.9 µm, EV-1106) using a standardized 21-minute gradient (Evosep 60 SPD method, 21 min) at 500 nL/min flow rate. Mass spectrometry data were acquired on a Q Exactive HF mass spectrometry system with nanospray Flex electrospray ionization source using both data-dependent and data-independent acquisition modes.

### Proteomics data analysis

Proteomics data are searched with ProLuCID^34^ and MSFragger^35^ search engines for DDA and DIA data using standard parameters. Specifically, 20 ppm precursor tolerance and 10 ppm fragment mass tolerance; standard trypsin digestion with up to 2 missed cleavage; Met and Pro oxidation, Ser, Thr, and Tyr phosphorylation were included as variable modifications according to MatrisomeDB^37,38^. Search results were filtered at 1% protein-level false discovery rate (FDR). A spectral library including the PTMs was constructed using DDA data from the same samples. The spectral library was then used to search DIA data and perform label-free quantitation. Sequence coverage was calculated and visualized using the Sequence Coverage Visualizer^36^.

### DNA extraction and analysis

DNeasy Blood and Tissue kit (#69504, QIAGEN) was used to extract DNA from samples according to the manufacturer’s instruction, except proteinase K digestion was done in the ATL buffer at 56 °C overnight. DNA was eluted in 100 μl of elution buffer. DNA concentration and purity (A 260/280) were measured by NanoDrop spectrophotometer (Thermo). The size of purified DNA was profiled with the genomic DNA ScreenTape assay (range: 200∼60000 bp) via the 4200 TapeStation system (Agilent).

### Construction of shotgun libraries

Construction of the DNA libraries was performed by the DNA Services Core at the Roy J. Carver Biotechnology Center, University of Illinois at Urbana-Champaign. Shotgun libraries were constructed using the Kapa Hyper Prep Sample Preparation Kit (Roche, Indianapolis, IN). For each sample, 150ng of DNA was run on 3 lanes of a 1% eGel (Thermo Fisher Scientific, Waltham, MA) and size-selected into two populations, 50-500nt (short), and 500+ (long). The sized samples were purified with MinElute columns (Qiagen, Hilden, Germany) and started in library prep with 10ng each for the ‘long’ samples and 8ng each for the ‘short’. Briefly, for the ‘long’ samples, DNA was sonicated on a Covaris M220 to a size of ∼ 400bp. The ‘short’ samples were not sonicated. All DNAs were then blunt-ended, 3’-end A-tailed, and ligated to unique dual-indexed adaptors. The adaptor-ligated DNA was amplified by PCR for 8 cycles with the Kapa HiFi polymerase (Roche, Indianapolis, IN). The final libraries were quantitated on Qubit and the average size determined on the AATI Fragment Analyzer (Agilent Technologies, Santa Clara, CA) and diluted to 5nM final concentration. The 5nM dilution was further quantitated by qPCR on a BioRad CFX Connect Real-Time System (Bio-Rad Laboratories, Inc. CA).

### DNA sequencing

Sequencing was performed by the DNA Services Core at the Roy J. Carver Biotechnology Center, University of Illinois at Urbana-Champaign. The final DNA library pool was sequenced on one lane of an Illumina NovaSeq 6000 SP flowcell v1.5 as paired-reads with 150nt length, to generate ∼1B reads in total, or ∼115M-155M read-pairs per sample. The run generated .bcl files which were converted into adaptor-trimmed demultiplexed fastq files using bcl2fastq v2.20 Conversion Software (Illumina, CA).

### DNA sequencing data analysis

The quality of the sequencing data was verified with FastQC. Bowtie2 (v2.5.1) using default settings to align each sample and sonication method to mm10 individually. The ‘long’ and ‘short’ sample bam files were merged using samtools (v1.6) to create one bam file for each respective sample. Bedgraphs were made with bedtools (v2.31.0) and were normalized by library size. To observe and visualize chromosomal sequencing depth on a macroscopic scale, we employed a sliding window over the normalized bedgraphs with a window size of 200kb and a step size of 20kb.

### Reconstitution of matrimeres *in vitro*

Genomic DNA was isolated from mouse D1 MSCs and sheared by sonication with a probe sonicator (VCX-130, Sonics & Materials) on ice at 30% amplitude for 10 s. Purified matrix proteins were added to autoclaved, double-filtered 0.1 M sodium acetate and acetic acid buffer solution (pH = 5.5) with the final concentration of 10 µg/ml, and stirred for 10 min at 4°C. The tested proteins include purified mouse FN (#IMSFBN, Innovative Research), purified mouse multimeric vitronectin (VTN; #IMSVTNMULTI, Innovative Research), and recombinant human FN III 12,13 N-GST (“HBDII”, which is derived from heparin-binding domain II of FN; #EUR120, Kerafast). Sheared DNA molecules were then added dropwise with continuous stirring at 400∼500 rpm with the final concentration of 10 µg/ml. In some cases, polyphosphate, long chain p700 (“polyP”; #EUI002, Kerafast) was used at 10 µg/ml instead of DNA. The reaction was continued for 8 h at 4°C. The solution was then centrifuged at 2,000g for 5 min to discard large aggregates, and nanoparticle tracking analysis was done afterward to determine particle size distribution and concentrations. To use reconstituted matrimeres for downstream analyses and experiments, the particles were loaded in 30% sucrose gradient for ultracentrifugation at 100,000g for 90 min at 4°C, followed by resuspending in double-filtered PBS. The average charge of reconstituted matrimeres was determined by measuring Zeta potential via Zetasizer (Malvern). The quantity of FN incorporated into reconstituted matrimeres was measured by first digesting them with DNase I (50 U/ml) for 1 h at 37 °C, followed by using the mouse FN enzyme-linked immunoassay (ELISA) kit (#E04552m, Lifeome) according to the manufacturer’s manual.

### Conjugation of reconstituted matrimeres with fluorescent dyes for *in vivo* experiments

FN-DNA matrimeres were freshly reconstituted and were dissolved in 2 mL Milli-Q water (2.0 x 10^10^ per mL). Then, 20 µL of 7.5 % Na_2_CO_3_ solution was added to maintain the pH at 6.5∼6.8. The solution was kept at 4 °C with the continuous stirring condition for 10 minutes to activate the amine. 40 µL of 10 mM Cy7-NHS Ester (#1115) or AZDye 488-NHS Ester (#1338) (both from Click Chemistry Tools) in DMSO was added to the solution dropwise in dark at 4 °C, and the reaction was continued for 12 h. The solution was dialyzed in a 14 kDa dialysis membrane (#D9277, Sigma) for 72 h against DI water to separate unconjugated dye molecules. The release of free dye molecules was studied by analyzing fluorescence intensity of the conjugated matrimeres (Cy7: ex: 753 nm, em: 775 nm; AZDye 488: ex: 494 nm, em: 517 nm) before and after dialysis, followed by resuspension in PBS after ultracentrifugation and particle counting by NanoSight.

### Tracking biodistribution of reconstituted matrimeres *in vivo*

4 h after treating mice with LPS (7.5 mg/kg) *i.p.*, Cy7-FN-DNA matrimeres (1.5 x 10^8^/20g) were delivered *i.t.* Mice were sacrificed at different time points (0.5, 8, 24, 72 and 216 h) for whole organ imaging by the IVIS Spectrum with the Living Image software (4.0, PerkinElmer) to quantify Cy7 fluorescence in radiance (photon.s^−1.^cm^−2.^sr^−1^) compared to untreated control mice. The images were acquired with the following parameters: 0.5 cm object height, low binning, 5 s exposure time, and 50% exposure power with an anti-glare mode.

### Flow cytometry analysis of immune cells to evaluate signals from reconstituted matrimeres after *in vivo* delivery

4 h after treating mice with LPS (7.5 mg/kg) *i.p.*, AZDye 488-FN-DNA matrimeres (1.5 x 10^8^/20g) were delivered *i.t.* Mice were perfused with PBS after 24 h to collect lung tissue. All the subsequent procedures were done in dark. To prepare single cell suspension, lung tissue was cut into small pieces and digested with collagenase (1 mg/ml; #C0130, Sigma) in DMEM with continuous shaking in water bath for 1 h at 37°C. The solution was then flushed by a syringe with a 16G needle (BD) until there was no visible tissue piece. The suspension was passed through a 40-µm filter, and remaining red blood cells were lysed with an ammonium-chloride-potassium (ACK) buffer (Thermo) for 1 min at RT, followed by centrifugation at 300g for 5 min to collect the cell pellet and washing 3 times with PBS. The single cell suspension was stained the following antibodies in PBS with 0.5% bovine serum albumin (BSA, Thermo) for 30 min at 4°C: CD45-APCCy7 (Clone 30-F11, 1:100), CD11b-PECy7 (Clone M1/70, 1:100), Ly6G-Pacific Blue (Clone 1A8, 1:100), CD11c-BV510 (Clone N418, 1:100), CD64-APC (Clone X54-5/7.1, 1:50), SiglecF-PE (Clone S17007L, 1:50), and MHCII-PE/Dazzle 594 (Clone M5/114.15.2, 1:50), all purchased from BioLegend. After staining, samples were washed 3 times with PBS/0.5% BSA, followed by fixation in 2% paraformaldehyde (Sigma) in PBS for 15 min. The samples were stored in PBS at 4°C for up to 1 week in dark before flow cytometry analysis by LSRFortessa (BD). The data were analyzed by the WEASEL software (2.7, WEHI). The following immune cell subpopulations were identified based on the previously published^84^ gating strategy (**Fig. S7**): neutrophils, lymphocytes, monocytes, monocyte-derived macrophages, tissue-resident alveolar macrophages, and interstitial macrophages. This was followed by mean intensity analysis of AZDye 488 in each immune subpopulation compared to untreated control mice.

### Investigating the spatial distribution of reconstituted matrimeres in lung tissue after *in vivo* delivery by two-photon microscopy

4 h after treating mice with LPS (7.5 mg/kg) *i.p.*, AZDye 488-FN-DNA matrimeres (1.5 x 10^8^/20g) were delivered *i.t*.., and the mice were kept alive for 24 or 216 h. 15 min before sacrificing the mice, 150 µL of 66.67 µg/mL of Alexa Fluor 647 anti-mouse CD31 Antibody (Clone MEC13.3, BioLegend) were administered *i.v.* Lung tissue was then harvested for two-photon imaging with the Ultima Multiphoton Microscope System (Bruker) to quantify the capillary network, collagen fibers, and labeled FN-DNA matrimeres. The Chameleon Ultra II Two-Photon laser operating at 80 MHz was used to excite lung tissue. Collagen fibers were excited at 860 nm and backward scattering of second harmonic generation was captured through a 430/24 nm bandpass filter. Vascular capillary images were acquired by exciting the Alexa Fluor 647-conjugated anti-mouse CD31 antibody signal at 810 nm and capturing the emission through a 708/75 bandpass filter. AZDye 488 labeled FN-DNA matrimeres were excited at 990 nm, and emission was recorded with a 525/50 bandpass filter. Z-stack images of all three signals were acquired from the same volume of parenchymal regions (defined as between 20 and 50 μm in depth from tissue surface with a field of view 149.1 μm x 149.1 μm) with 1 μm interval via Prairie View software (5.4, Bruker) using an Olympus XLUMPlanFL 20x/1.00NA objective.

### Three-dimensional two-photon image analysis

Imaris (9.3.1, Bitplane) was used to perform 3D reconstruction of two-photon images from each stack. Voxels were generated in green, red, and blue signals to represent AZDye 488 FN-DNA matrimeres, Alexa Flour 647-CD31 blood vessels, and collagen fibers, respectively. Thresholding values were set with variations less than 20% across all the images from different experiments. The total voxels above the threshold were calculated to quantify the surface area of each cluster in each channel per field of view. The green particles with greater than 150 μm^2^ in surface area (CED ∼ 7 μm) were considered ‘cell-like’ particles, while the particles with less than 30 μm^2^ in surface area (CED ∼ 3 μm) were considered ‘small’ clusters. The contact surface areas between FN-DNA matrimeres and blood vessels (yellow), and between FN-DNA matrimeres and collagen fibers (cyan) were determined by the built-in algorithm (XTension) in Imaris.

### Lung function measurement *in vivo*

Mice were anesthetized with ketamine/xylazine (50/5 mg/kg) *i.p.* and intubated. The mice were then connected to the FlexiVent (SCIREQ) system, and the lungs were inflated up to 30 cmH_2_O in pressure. The forced oscillation technique was used to derive the following functional parameters: inspiratory capacity (IC, mL), tissue damping (G, cmH_2_O^s^mL^-1^), respiratory system compliance (Crs, mL cmH_2_O^−1^), and respiratory system elastance (Ers, cmH2O mL^−1^). The measurements were repeated 3 times per mouse to obtain average values for each parameter. Mice were allowed to recover in a heated recovery chamber after the measurements.

### Cutaneous reverse passive Arthus (rpA) reaction assay

Balb/cJ mice at the age of 8∼10 weeks were anesthetized by ketamine/xylazine (100/16 mg/kg) *i.p.*, followed by *i.v.* injection of BSA (75 μg/g) as well as indicated treatments in 100 μl sterile 0.9% NaCl. To evaluate vascular permeability, EBA was *i.v.* injected simultaneously. The medial surface of the back was shaved for each mouse recipient. After 5 min, mice were intradermally injected with rabbit anti-BSA antibody (6 mg/ml, MP Biomedicals) in 25 μl of PBS, while control sites received 25 μl of PBS without the antibody in parallel. Evans blue extravasation in the skin samples was measured (absorbance: 620 nm) after 4 h, and the hemoglobin contents were quantified after 8 h by the colorimetric assay kit (absorbance: 550 nm; with BD Pharm Lyse). Each value was subtracted from the internal control without the anti-BSA antibody.

### Statistical Analysis

Statistics were performed as described in figure captions. All statistical analyses were performed using GraphPad Prism version 10.1.0. Unless otherwise noted, statistical comparisons were made by one-way ANOVA followed by Tukey’s multiple comparisons test when standard deviations did not vary between test groups, and by one-way Welch ANOVA followed by Dunnett T3 multiple comparisons test when standard deviations were variable. A *p*-value less than 0.05 established statistical significance.

## Acknowledgments

This work made use of instruments in the Flow Cytometry Core, Fluorescence Imaging Core, Genomics Research Core, and Electron Microscopy Core in the Research Resources Center at UIC. We acknowledge the Cardiovascular Research Core at the UIC Research Resources Center (Ayman Isbatan and Jiwang Chen) for providing technical expertise in lung function measurements and the DNA Services Core at the Roy J. Carver Biotechnology Center, University of Illinois at Urbana-Champaign for performing DNA sequencing analysis. This work was supported by the National Institutes of Health Grants, R01EB034507, R01HL141255, R01GM141147 (J-W.S.); R35GM133416 (Y.G.); R35HL150797 (X.D.); P01HL151327, R01HL152515 (J.R.); R01HL084153 (D.M.); and the National Science Foundation Grant CAREER 2143857 (J-W.S.).

## Author contributions

Conceptualization, K.D., J-W.S.; Data curation, K.D., J-W.S.; Formal analysis, K.D., J.E., C.L., M.S., J-W.S.; Funding acquisition, D.M., J.R., X.D., Y.G., J-W.S.; Investigation, K.D., I.Q., S.O., J.E., C.W., M.S., C.L., A.R, I.S.C., S.S., P.T., Y.G., J-W.S.; Methodology, K.D., J.E., C.W., M.S., C.L., I.S.C., P.T., J.R., X.D., Y.G., and J-W.S.; Project administration, J-W.S.; Resources, D.M., J.R., X.D., Y.G., and J-W.S.; Software, M.S., C.L., J-W.S.; Supervision, J-W.S.; Validation, I.Q., S.O., A.R., and J-W.S.; Visualization, M.S., C.L., J-W.S.; Writing original draft, J-W.S.; Editing original draft, K.D., J.E., M.S., D.M., J.R., X.D., Y.G., J-W.S.

## Inclusion & ethics statement

All contributors to this research meet the authorship criteria specified by Nature Portfolio journals and have been acknowledged as authors. Their involvement was crucial for the study’s design and execution. Collaborators established their roles and responsibilities before the research started. The study faced no significant constraints or prohibitions in the researchers’ environment, ensuring it does not lead to stigmatization, incrimination, discrimination, or personal risk for the participants.

## Data availability

The data that support the plots within this paper and other findings of this study are available from the corresponding author upon reasonable request.

## Code availability

The codes used to analyze the data in this study are available from the corresponding author upon reasonable request.

