## Supplementary Information for "Matrimeres are systemic nanoscale mediators of tissue integrity and function"

**Supplementary Text**

**1. Calculation of different nanoscale subpopulations secreted by cells**

FN: Fibronectin

CD63: Tetraspanin (membrane protein)

TX: Triton-X (+: sensitive, -: resistant)

Each set of the values (*n* = 3 independent experiments) from Fig. 1A was used to calculate 8 subpopulations in terms of FN, CD63 and TX. The values in the following calculations are from one of the replicate sets. These values were used to construct Fig. 1A, iii.

1. The remaining fraction of nanoscale mediators after depleting with FN and CD63 antibodies and treating with Triton-X is expressed as follows.

$$\begin{aligned}\boldsymbol{P(FN^- \cap CD63^- \cap TX^-)} &= P(FN^- \cap CD63^-) \times P(TX^- \mid FN^- \cap CD63^-) \\ &= (1 - P(FN^+ \cup CD63^+)) \times (1 - P(TX^+ \mid FN^- \cap CD63^-)) \\ &= (1 - 0.9597)\ (Fig. 1A, i) \times (1 - 0.2278)\ (Fig. 1A, ii) \\ &= \mathbf{0.0311}\ (Eq. 1.1)\end{aligned}$$

2. From #1, the negative sign of each marker can be switched to the positive sign one at a time to calculate the following subpopulations.

(a) $P(FN^+ \cap CD63^- \cap TX^-) = P(CD63^- \cap TX^-) - P(FN^- \cap CD63^- \cap TX^-)$

$$\begin{aligned}P(CD63^- \cap TX^-) &= P(CD63^-) \times P(TX^- \mid CD63^-) \\ &= (1 - P(CD63^+)) \times (1 - P(TX^+ \mid CD63^-)) \\ &= (1 - 0.52258)\ (Fig. 1A, i) \times (1 - 0.0755)\ (Fig. 1A, ii) = 0.4414\ (Eq. 1.2)\end{aligned}$$

$$\boldsymbol{P(FN^+ \cap CD63^- \cap TX^-)} = 0.4414 - 0.0311\ (Eq. 1.1) = \mathbf{0.4103}\ (Eq. 1.3)$$

(b) $P(FN^- \cap CD63^+ \cap TX^-) = P(FN^- \cap TX^-) - P(FN^- \cap CD63^- \cap TX^-)$

$$P(FN^- \cap TX^-) = P(FN^-) \times P(TX^- \mid FN^-)$$
$$= (1 - P(FN^+)) \times (1 - P(TX^+ \mid FN^-))$$
$$= (1 - 0.4642)\ (Fig. 1A, i) \times (1 - 0.8424)\ (Fig. 1A, ii) = 0.0844\ (Eq. 1.4)$$

$$\boldsymbol{P(FN^- \cap CD63^+ \cap\ TX^-)} = 0.0844 - 0.0311\ (Eq. 1.1) = \mathbf{0.0533}\ (Eq. 1.5)$$

(c) $\boldsymbol{P(FN^- \cap CD63^- \cap\ TX^+)} = \ P(FN^- \cap CD63^-) - P(FN^- \cap CD63^- \cap\ TX^-)$
$$= \ 1 - P(FN^+ \cup CD63^+)\ - P(FN^- \cap CD63^- \cap\ TX^-)$$
$$= \ 1 - 0.9597\ (Fig. 1A, i) - \ 0.0311\ (Eq. 1.1) = \ \mathbf{0.0092}\ (Eq. 1.6)$$

3. From #2, the negative sign of one marker can be switched to the + sign to calculate the following subpopulations.

(a) $P(FN^+ \cap CD63^+ \cap\ TX^-) = \ P(FN^+ \cap TX^-) - P(FN^+ \cap CD63^- \cap\ TX^-)$

$$P(FN^+ \cap TX^-) = \ P(TX^-) - \ P(FN^- \cap TX^-)$$
$$= (1 - P(TX^+)) - \ P(FN^- \cap TX^-)$$
$$= (1 - 0.5016)\ (Fig. 1A, ii) - \ 0.0844\ (Eq. 1.4) = 0.414\ (Eq. 1.7)$$

$$\boldsymbol{P(FN^+ \cap CD63^+ \cap\ TX^-)} = 0.414 - 0.4103\ (Eq. 1.3) = \mathbf{0.0037}\ (Eq. 1.8)$$

(b) $P(FN^- \cap CD63^+ \cap\ TX^+) = \ P(FN^- \cap CD63^+) - P(FN^- \cap CD63^+ \cap\ TX^-)$

$$P(FN^- \cap CD63^+) = \ P(FN^-) - \ P(FN^- \cap CD63^-)$$
$$= (1 - P(FN^+)) - (1 - P(FN^+ \cup CD63^+))$$
$$= (1 - 0.4642)\ (Fig. 1A, i) - (1 - 0.9597)\ (Fig. 1A, i) = 0.4955\ (Eq. 1.9)$$

$$\boldsymbol{P(FN^- \cap CD63^+ \cap\ TX^+)} = \ 0.4955 - 0.0533\ (Eq. 1.5) = \mathbf{0.4422}\ (Eq. 1.10)$$

(c) $P(FN^+ \cap CD63^- \cap\ TX^+) = \ P(CD63^- \cap TX^+) - P(FN^- \cap CD63^- \cap\ TX^+)$

$$P(CD63^- \cap TX^+) = \ P(CD63^-) - \ P(CD63^- \cap TX^-)$$

$$= (1 - P(CD63^{+})) - P(CD63^{-} \cap TX^{-})$$
$$= (1 - 0.5226)\ (Fig.\,1A, i) - \ 0.4414\ (Eq.\,1.2) = 0.036\ (Eq.\,1.11)$$

$$\boldsymbol{P}(\boldsymbol{FN}^{+} \cap \boldsymbol{CD63}^{-} \cap \ \boldsymbol{TX}^{+}) = 0.036 - 0.0092\ (Eq.\,1.6) = \mathbf{0.0268}\ (Eq.\,1.12)$$

4. Since the total fraction of the subpopulations is 1, the fraction of the subpopulation that is positive for both FN and CD63 and is sensitive to TX can be calculated from #1-3 as follows.

$\boldsymbol{P}(\boldsymbol{FN}^{+} \cap \boldsymbol{CD63}^{+} \cap \ \boldsymbol{TX}^{+}) = 1 \ - \big(P(FN^{-} \cap CD63^{-} \cap \ TX^{-}) + \ P(FN^{+} \cap CD63^{-} \cap \ TX^{-}) + P(FN^{-} \cap CD63^{+} \cap \ TX^{-}) + \ P(FN^{-} \cap CD63^{-} \cap \ TX^{+}) + \ P(FN^{+} \cap CD63^{+} \cap \ TX^{-}) + \ P(FN^{-} \cap CD63^{+} \cap \ TX^{+}) + \ P(FN^{+} \cap CD63^{-} \cap \ TX^{+})\big) = 1 - (0.0311 + 0.4103 + 0.0533 + 0.0092 + 0.0037 + 0.4422 + 0.0268) = \mathbf{0.0234}\ (Eq.\,1.13)$

**2. Calculation of the number of FN and DNA molecules per reconstituted FN-DNA particle**

The number of FN-DNA particles produced from 10 μg/ml FN and 10 μg/ml DNA at pH ~ 5.5 is 1.5 x $10^{10}$/ml (**Fig. 4A, ii**). Dividing 1.5 x $10^{10}$ by the Avogadro's number (6.0 x $10^{23}$/mol) yields 2.5 x $10^{-14}$ mol of FN-DNA particles.

4.6 μg/ml FN is incorporated into 1.5 x $10^{10}$/ml FN-DNA particles (**Fig. 4A, iv**). Since the molecular weight of monomeric FN is ~220 kDa but FN normally exists as a dimer (i.e., 440,000 g/mol), this means 1.05 x $10^{-11}$ mol of FN is incorporated into 2.5 x $10^{-14}$ mol of FN-DNA particles. Thus, ~420 molecules of FN dimers or ~840 FN monomers are incorporated into a single FN-DNA particle.

5 μg/ml genomic double-stranded DNA is incorporated into 1.5 x $10^{10}$/ml FN-DNA particles (**Fig. 4A, v**). The average molecular weight of a single DNA base pair is 650 g/mol, which means 7.7 x $10^{-9}$ mol of DNA base pairs is incorporated into 2.5 x $10^{-14}$ mol of FN-DNA particles. Thus, ~307,700 DNA base pairs are incorporated into a single FN-DNA particle.

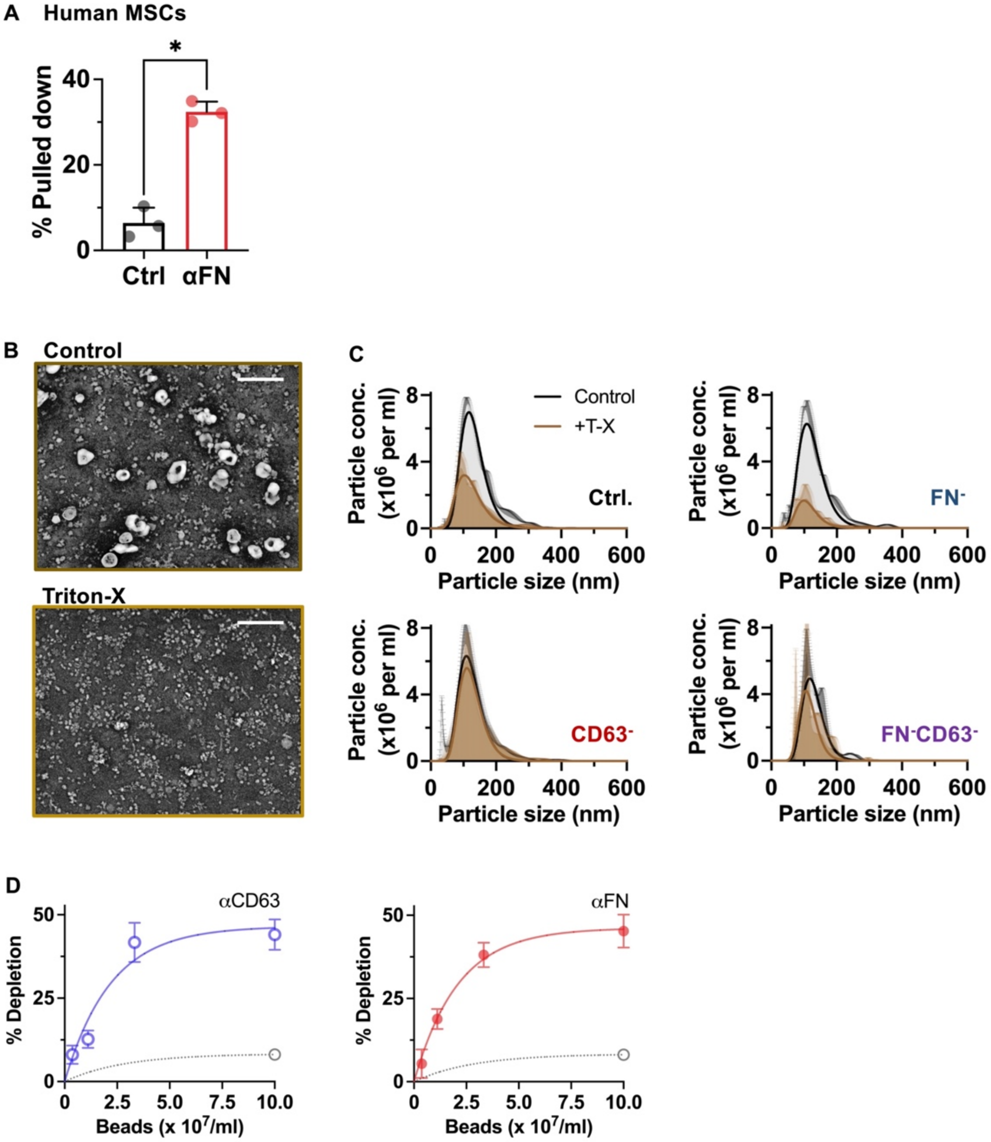


**Figure S1. Characterization of cell-secreted small nanostructures. (A)** Percentage of $FN^+$ particles secreted from human primary bone marrow MSCs. $n = 3$ donors, $*p < 0.05$, via unpaired T-test. **(B)** Representative images of mouse bone marrow D1 MSC-secreted small (< 200 nm) nanostructures before and after treatment with 0.1% (v/v) Triton-X. Scale bar = 400 nm. **(C)** Hydrodynamic sizes of mouse MSC-secreted small nanostructures. The average data points were fitted to log-normal distribution curves. $n = 3$ experiments. **(D)** Mouse MSC-secreted small nanostructures (~5.0 x $10^8$) were mixed with a varying number of (Left) CD63 or (Right) FN antibody-coated magnetic beads, and the percentage of the nanostructures pulled down (or depleted) by the beads was calculated. The data points were fitted to one-phase association curves. Half-maximum bead number and plateau for CD63 antibody: 1.5 x $10^7$, 46.5%; FN antibody: 1.4 x $10^7$, 46.2%, from $n = 3$ experiments. The dotted lines and a grey data point show the background percentage of pull-down with biotin-coated beads.

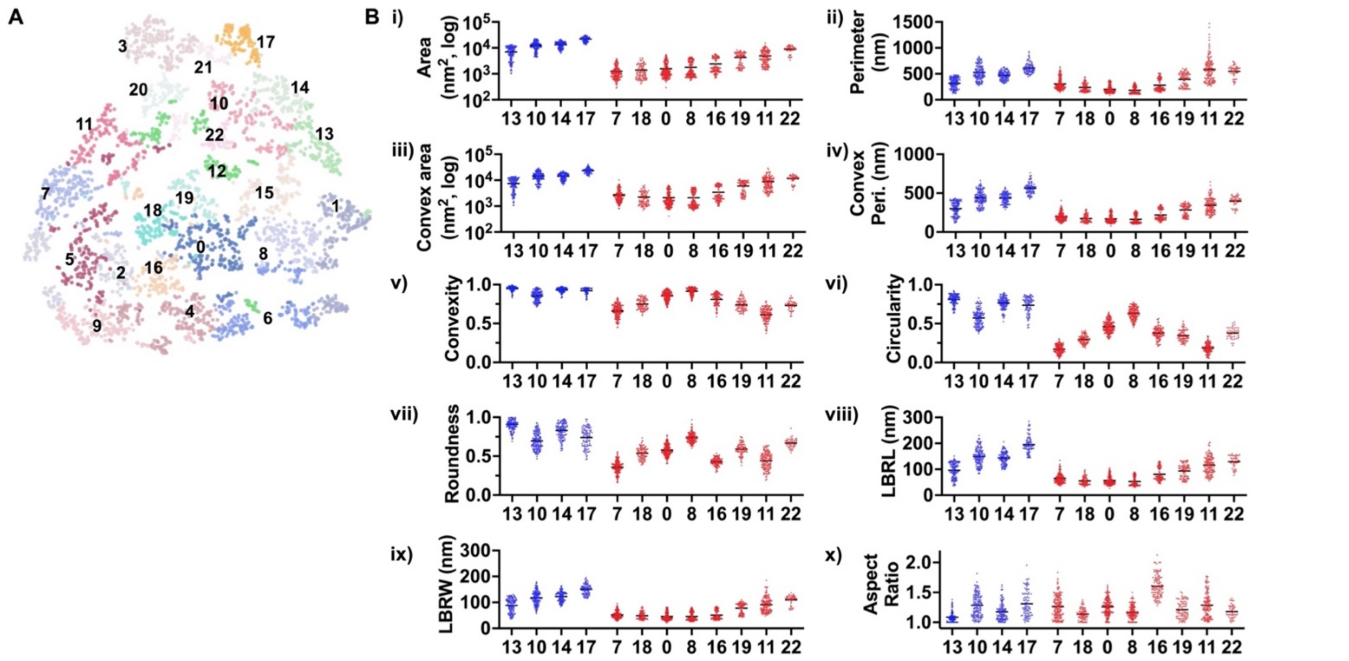


**Figure S2. Multiparameter morphological analysis of cell-secreted nanostructures. (A)** T-distributed stochastic neighbor embedding (t-SNE) plot showing different clusters of mouse bone marrow D1 MSC-secreted small nanostructures based on morphological analysis of TEM images from non-depleted (ctrl., *n* = 1581), FN-depleted (*n* = 895), and CD63-depleted (*n* = 679) samples. The clusters and corresponding numbers were assigned by using the Leiden algorithm, and are shown in different colors. **(B)** Quantification of morphological parameters for each cluster. (i) Area. (ii) Perimeter. (iii) Convex area. (iv) Convex perimeter. (v) Convexity. (vi) Circularity. (vii) Roundness. (viii) Least bounding rectangle length (LBRL). (ix) Least bounding rectangle width (LBRW). (x) Aspect ratio. The blue color shows the clusters that are more represented by FN$^-$ than CD63$^-$ nanostructures, while the red color shows the clusters that are more represented by CD63$^-$ than FN$^-$ nanostructures. *n* = 47-198 for each group.

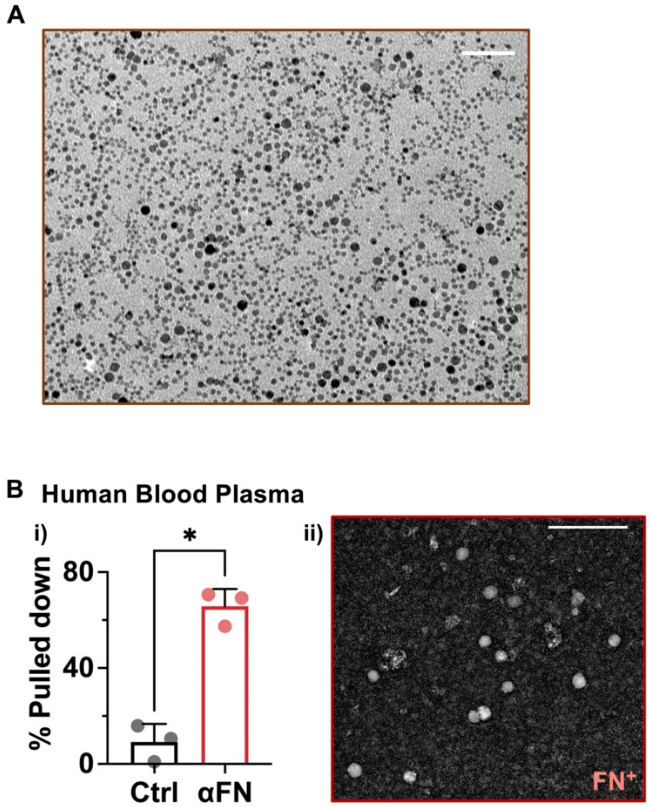


**Figure S3. Imaging of FN$^+$ matrimeres after positive selection with antibody-conjugated magnetic nanoparticles. (A)** Representative TEM image showing synthesized magnetic nanoparticles for positive selection of a specific (CD63$^+$ or FN$^+$) nanostructure subpopulation. Scale bar = 50 nm. **(B)** FN$^+$ matrimeres in human blood plasma. (i) Percentage of FN$^+$ matrimeres. $n$ = 3 donors, $*p < 0.05$, via unpaired T-test. (ii) Representative TEM image showing a group of FN$^+$ matrimeres after positive selection. Scale bar = 200 nm.

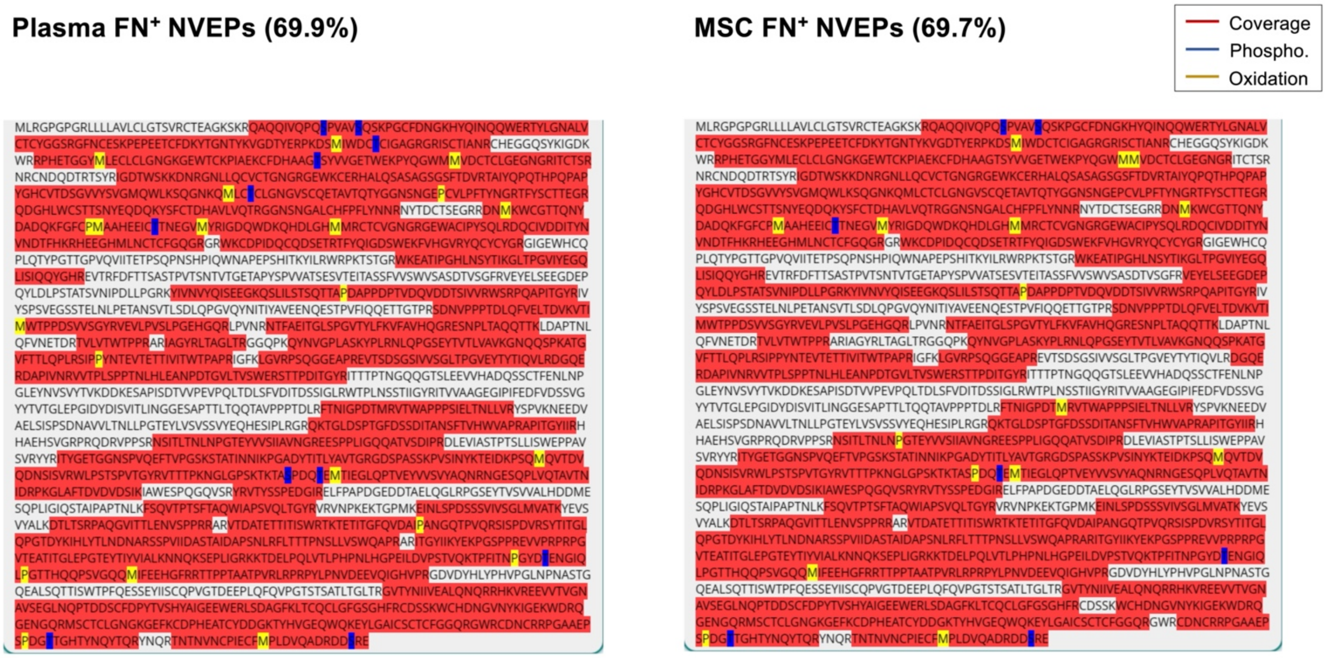


**Figure S4. Detailed sequence coverage of Fn1 from MSC *vs.* plasma FN$^{+}$ NVEPs.** Red denotes detected regions; histograms above denote peptide overlap density; blue and yellow denote the phosphorylation and oxidation sites detected, respectively.

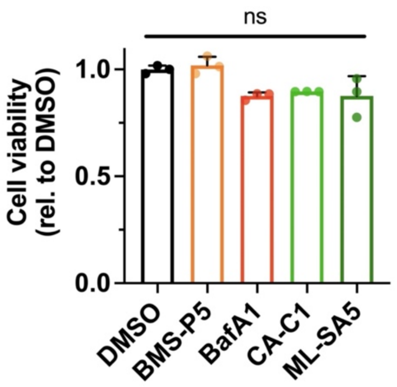


**Figure S5. Effects of pharmacological agents on cell viability.** An equal number of mouse D1 MSCs were seeded and cultured in the presence of DMSO or one of the tested drugs for 24 h. Cells were then detached to count the viable cell number by trypan blue exclusion, which was normalized to the mean cell number of the DMSO group.

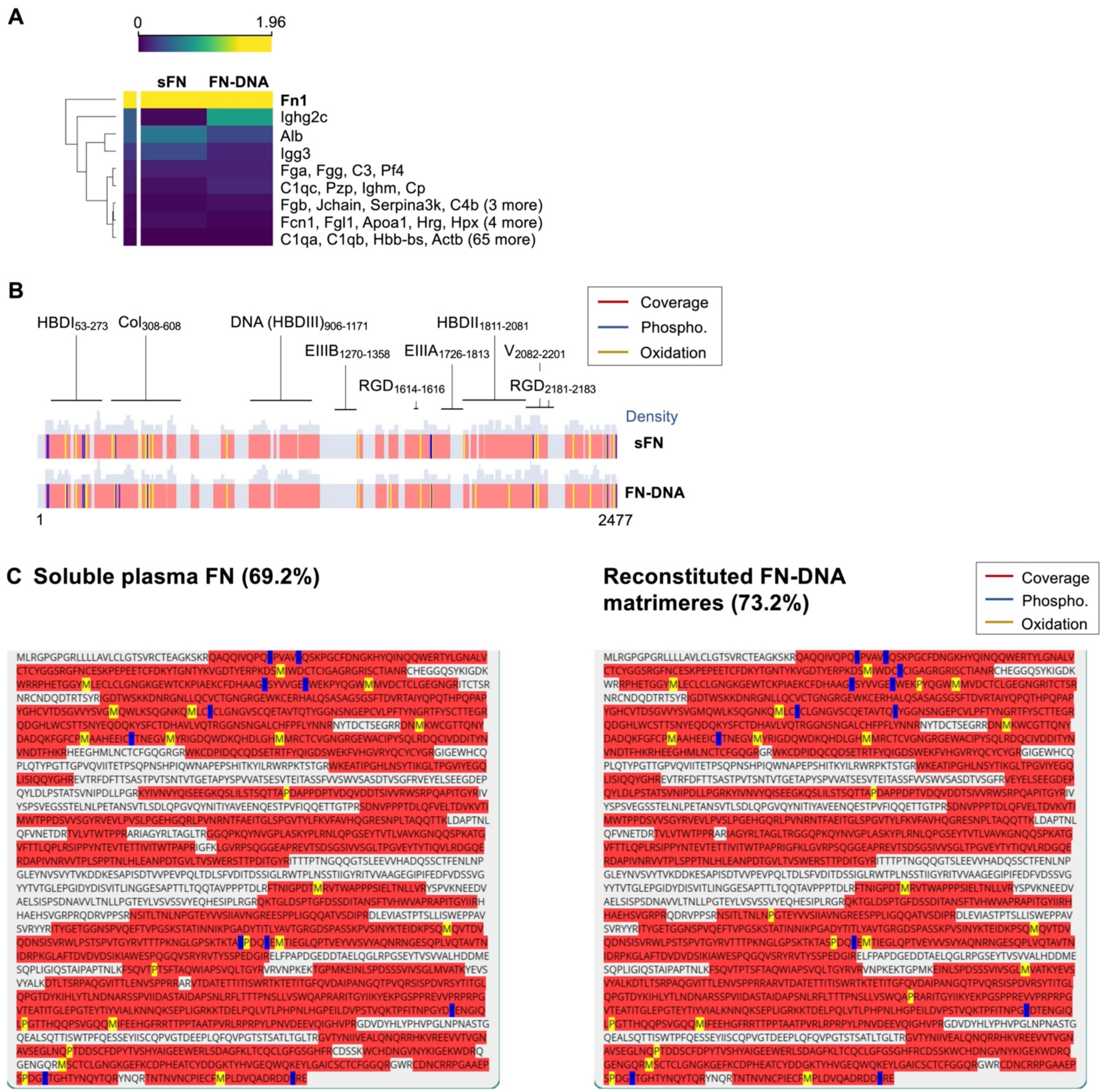


**Figure S6.** Proteomic analysis of soluble blood plasma FN (sFN) before and after incorporation into FN-DNA matrimeres. **(A)** Proteome coverage comparison; label-free protein quantitation based on log-transformed intensity. **(B)** Fn1 sequence coverage comparison. **(C)** Detailed sequence coverage of Fn1. Red denotes detected regions; histograms above denote peptide overlap density; blue and yellow denote the phosphorylation and oxidation sites detected, respectively. Pooled from *n* = 2 biological replicates.

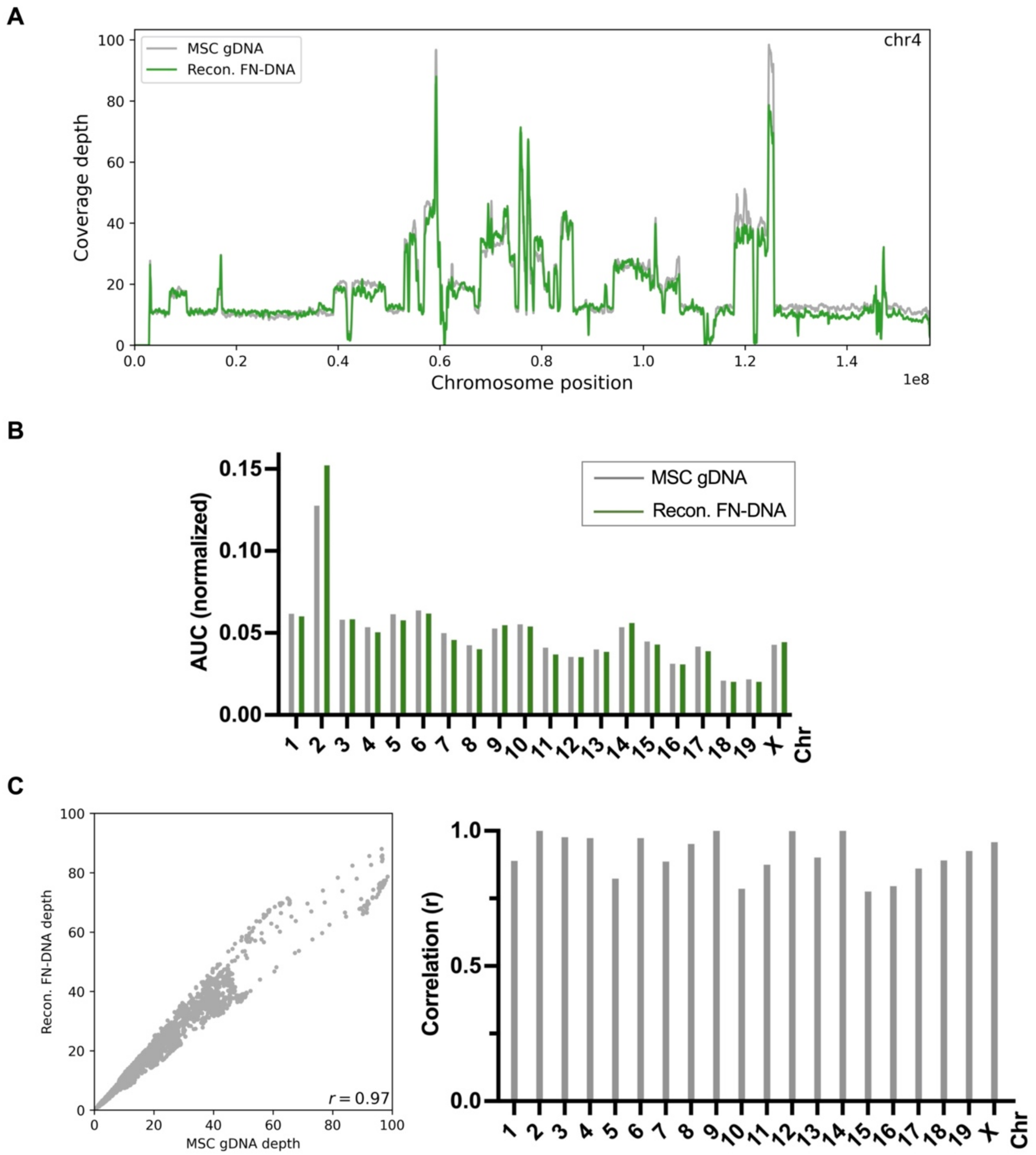


**Figure S7. Comparison of genome-wide chromosomal coverage between genomic DNA of mouse MSCs and sheared genomic DNA assembled into reconstituted FN-DNA matrimeres. (A)** Representative graph showing the sequence coverage of chromosome 4. **(B)** Area under the curve (AUC) of coverage depth for each chromosome. **(C)** Correlation analysis of coverage depth values between DNA of reconstituted FN-DNA matrimeres and genomic DNA of MSCs. (Left) Representative graph of chromosome 4. (Right) Pearson's correlation coefficient (*r*) values of each chromosome. One biological replicate for each group.

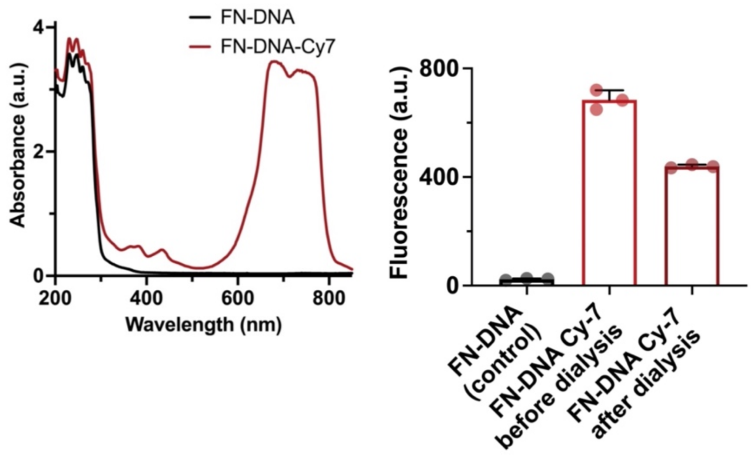


**Figure S8. Characterization of Cy7-conjugated reconstituted FN-DNA matrimeres.** (Left) Representative absorption spectra of Cy7-conjugated *vs.* unconjugated FN-DNA matrimeres. (Right) Mean fluorescence intensity of FN-DNA matrimeres after conjugation with Cy7 before and after dialysis.

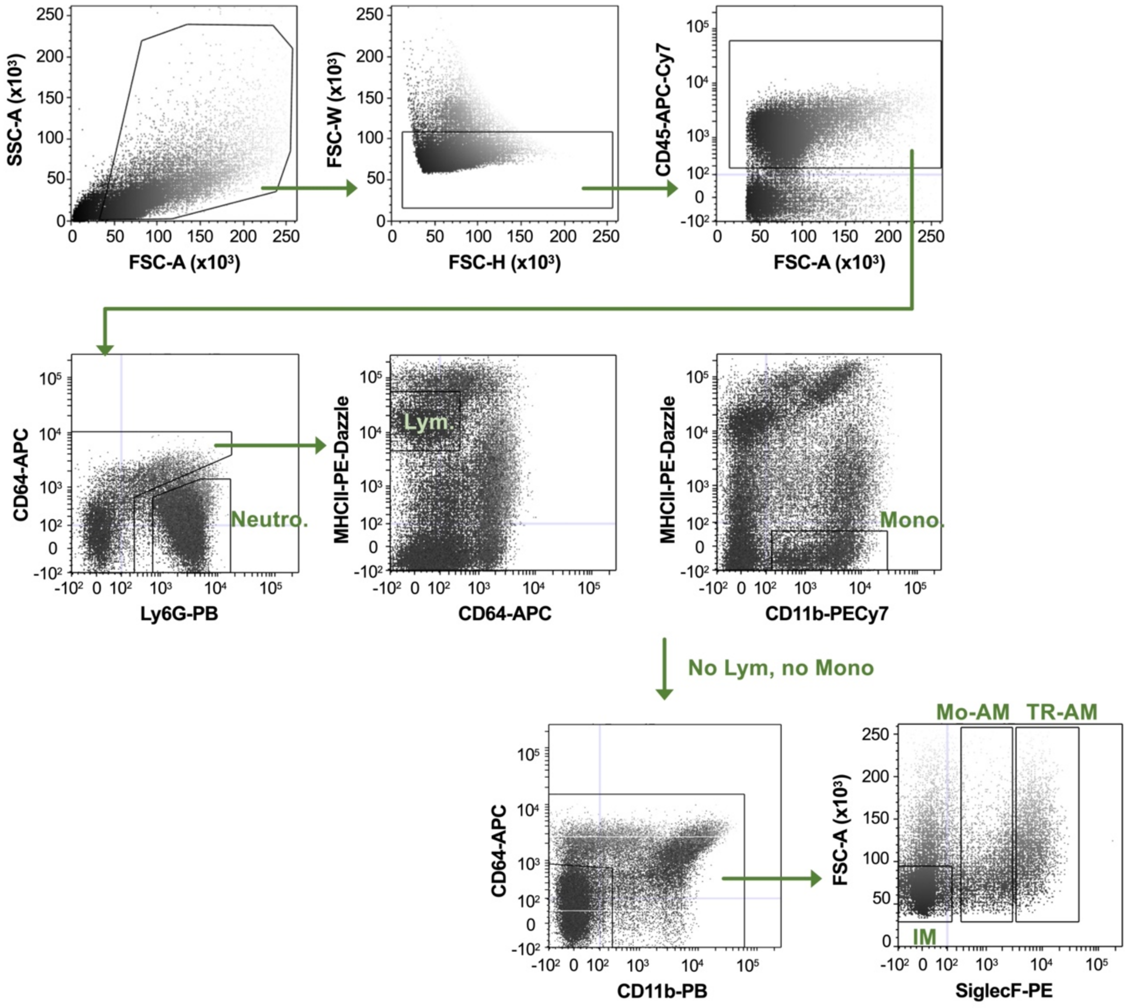


**Figure S9. Gating strategy to identify immune cell subpopulations.** After isolating single cell suspension from mouse lung tissue by enzymatic digestion, immune cells were identified by $CD45^+$ gating, along with exclusion of aggregates. A sequential gating strategy was first used to identify neutrophils (Neutro., $Ly6G^+$), lymphocytes (Lym., $CD64^-MHCII^+$), and monocytes ($CD11b^+MHCII^-$). Macrophage subpopulations were identified after excluding lymphocyte and monocyte populations, followed by excluding $CD64^-CD11b^-$ populations. $SiglecF^-$: interstitial macrophages (IM), monocyte-derived alveolar macrophages (Mo-AM, $SiglecF^{lo}$), and tissue-resident alveolar macrophages (TR-AM, $SiglecF^{hi}$).
